# Bright and tunable far-red chemigenetic indicators

**DOI:** 10.1101/2020.01.08.898783

**Authors:** Claire Deo, Ahmed S. Abdelfattah, Hersh K. Bhargava, Adam J. Berro, Natalie Falco, Benjamien Moeyaert, Mariam Chupanova, Luke D. Lavis, Eric R. Schreiter

**Author notes:** These authors contributed equally: Claire Deo, Ahmed S. Abdelfattah. These authors contributed equally: Luke D. Lavis, Eric R. Schreiter.

## Abstract

Functional imaging using fluorescent indicators has revolutionized biology but additional sensor scaffolds are needed to access properties such as bright, far-red emission. We introduce a new platform for ‘chemigenetic’ fluorescent indicators, utilizing the self-labeling HaloTag protein conjugated to environmentally sensitive synthetic fluorophores. This approach affords bright, far-red calcium and voltage sensors with highly tunable photophysical and chemical properties, which can reliably detect single action potentials in neurons.

---

Fluorescent sensors enable noninvasive measurement of cellular function. Although this field began with small-molecule fluorescent dyes, functional imaging in biology rapidly switched to protein-based sensors after the discovery of green fluorescent protein (GFP). Protein-based indicators allow cell-specific expression and take advantage of evolved molecular recognition motifs found in nature. In particular, genetically encoded calcium and voltage indicators (GECIs and GEVIs) have become powerful tools for monitoring neuronal activity in living systems.^1, 2^ Commonly used GECIs and GEVIs, such as GCaMP^3, 4^ and ASAP^5^, rely on the use of a circularly permuted (cp) GFP fused to sensor protein domains like calmodulin (CaM) or a voltage sensitive domain (VSD). A conformational change in the sensor domain alters the environment around the GFP chromophore, resulting in a fluorescence change (Fig. 1a). Although this approach has led to robust sensors excitable with shorter wavelengths (<550 nm), analogous sensors based on red-shifted fluorescent proteins have proven much more difficult to optimize.^6, 7^ Here, we demonstrate a new hybrid^8, 9^ small-molecule:protein (‘chemigenetic’) sensor scaffold based on the self-labeling HaloTag protein^10, 11^ and fluorogenic rhodamine dyes^12–15^. These chemigenetic sensors combine the genetic targetability and exquisite molecular recognition of proteins with the improved photophysical properties of synthetic fluorophores. We showcase the advantages of this system by creating new bright, far-red calcium and voltage sensors with tunable photophysical and chemical properties that are capable of robust reporting of individual action potentials in single-trial recordings from neurons.

**Fig. 1.**
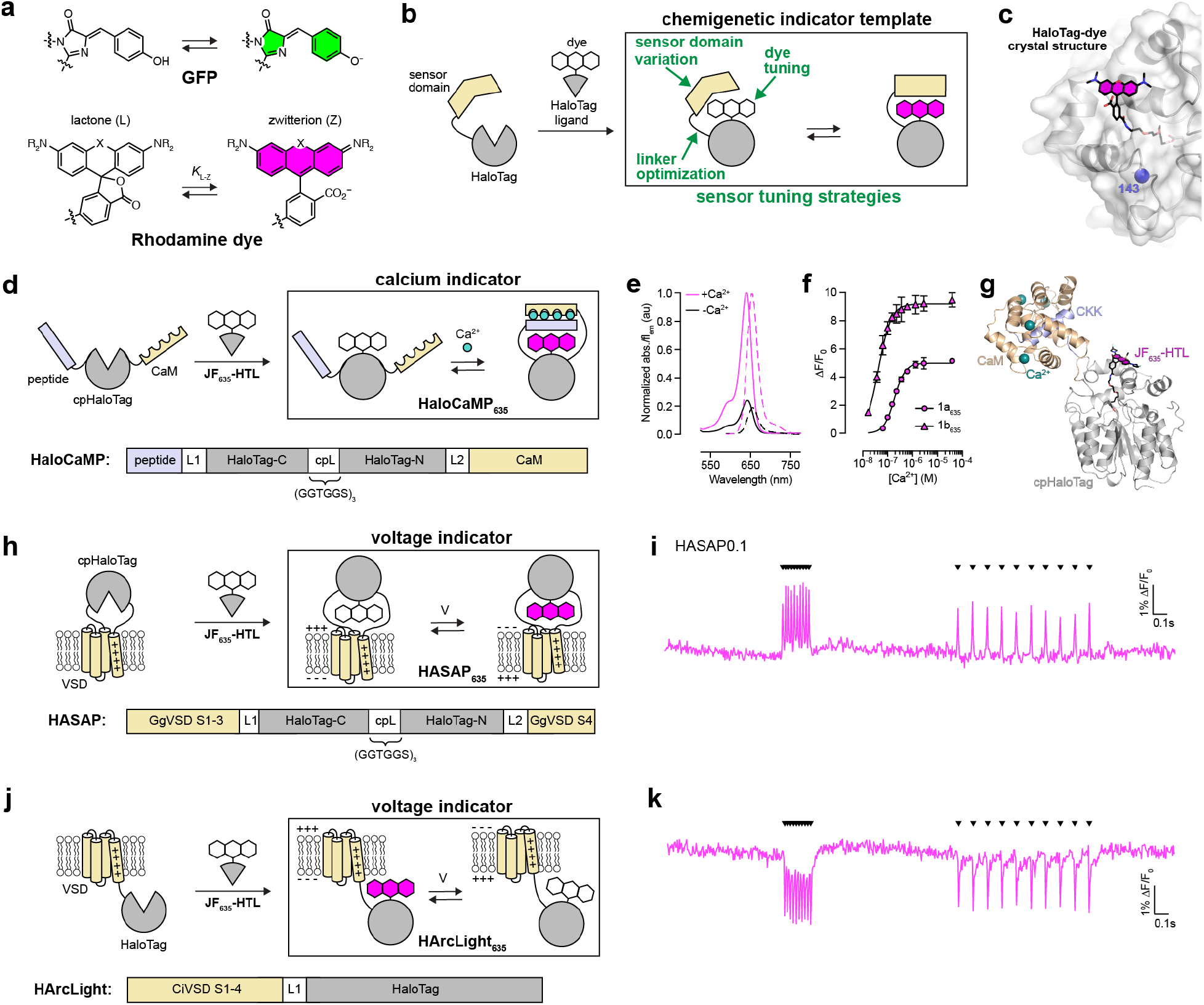
Strategy for engineering chemigenetic indicators. **a**, Non-fluorescent/fluorescent equilibria of the chromophore of GFP (top) and a rhodamine dye (bottom). **b**, Schematic representation of our strategy for chemigenetic sensor engineering. A sensor protein domain is fused to HaloTag such that after binding of a dye-ligand, conformational changes within the sensor protein domain alter the environment and fluorescence output of the bound dye. **c**, Crystal structure of HaloTag bound to TMR-HTL. A cartoon and surface representation of the HaloTag protein are shown in grey, the bound TMR-HTL is black lines with the xanthene ring system of the rhodamine dye filled in magenta. Position 143 of HaloTag, where it was circularly permuted to create sensors, is shown as a blue sphere. **d**, Schematic of the chemigenetic calcium indicator HaloCaMP, showing domain organization (top) and primary structure (bottom). **e**, Normalized absorption and fluorescence spectra of HaloCaMP1a_635_. **f**, Calcium titration of HaloCaMP1a_635_ and HaloCaMP1b_635_. Mean and s.d. for *n* = 2 independent titrations. **g**, Crystal structure of Ca^2+^-saturated HaloCaMP1b_635_. **h**, Schematic of the chemigenetic voltage indicator HASAP, showing domain organization (top) and primary structure (bottom). **i**, Fluorescence response of HASAP0.1 in cultured hippocampal neurons labeled with **JF**_**635**_**-HTL** to a train of field stimuli eliciting 10 action potentials at 50 Hz followed by 10 action potentials at 10 Hz. Field stimulus timing shown with inverted triangles. **j**, Schematic of the chemigenetic voltage indicator HArclight, showing domain organization (top) and primary structure (bottom). **k**, Fluorescence response of HArcLight1 in cultured hippocampal neurons labeled with **JF**_**635**_**-HTL** to a train of field stimuli eliciting 10 action potentials at 50 Hz followed by 10 action potentials at 10 Hz. Field stimulus timing shown with inverted triangles.

The impetus for our chemigenetic scaffold described here stems from the recent development of fluorogenic ligands and stains based on rhodamine dyes. Rhodamine derivatives exist in equilibrium between a closed, non-fluorescent lactone (L) form and an open, fluorescent zwitterionic (Z) form (Fig. 1a). This L–Z equilibrium constant (*K*_L-Z_)—and therefore the fluorescence intensity—is strongly environment dependent.^16^ Although classic rhodamine dyes such as tetramethylrhodamine (TMR) strongly favor the zwitterionic form, other rhodamine analogs such as the far-red Si-rhodamine Janelia Fluor 635 (**JF**_**635**_) exist predominantly in the lactone form in aqueous solution.^15^ Upon binding the HaloTag protein, however, the corresponding **JF**_**635**_–HaloTag ligand shows >100-fold increase in absorbance and fluorescence due to the change in local environment. Further examination of the spectral properties of **JF**_**635**_-HaloTag conjugates revealed that binding did not fully shift the equilibrium to the zwitterionic form; the absorptivity of the **JF**_**635**_–HaloTag conjugate (ε = 81,000 M^−1^cm^−1^) was lower than the maximal value of **JF**_**635**_ determined in acidic solution (ε_max_ = 167,000 M^−1^cm^−1^)^15^. Based on this observation, we reasoned that the **JF**_**635**_–HaloTag pair could be used to create chemigenetic fluorescent biosensors where conformational changes within a protein sensor domain could change the dye environment on the HaloTag, resulting in a shift in the L–Z equilibrium and modulation of fluorescence (Fig. 1b).

To guide our design of new biosensors, we solved a crystal structure of HaloTag labeled with the TMR-HaloTag ligand (Fig. 1c, Supplementary Table 1). The bound dye appears in the open, zwitterionic form, and adopts multiple conformations associated with the surface of HaloTag. This suggests that both conformational restrictions imposed by tethering to the protein as well as interactions with specific residues at the protein surface are responsible for the change in fluorophore environment that shifts the L–Z equilibrium of dyes such as **JF**_**635**_ towards the fluorescent zwitterionic form upon protein binding. Using this structure, we identified and tested possible sites for circular permutation to create new N- and C-termini in close spatial proximity to the bound fluorophore (Supplementary Fig. 1), analogous to the circular permutation of GFP within beta strand seven near its chromophore^17^. We found that circular permutation within the loop around position 143 of HaloTag (cpHaloTag) was well-tolerated. We tested the generality of this approach by designing chemigenetic calcium and voltage indicators inspired by the existing single FP-based indicators: GCaMP^3, 4^, ASAP^5^, and Arclight^18^ (Fig. 1d-k). Our chemigenetic sensor scaffold can be modulated by changing the sensor protein domain, by mutagenesis of the linkers connecting the sensor and HaloTag domains, and by changing the synthetic fluorophore (Fig. 1b).

We created Ca^2+^ sensors by appending CaM to the new cpHaloTag C-terminus and the CaM-binding peptides from myosin light chain kinase (MLCK) or CaM-dependent kinase kinase (CKK) to the N-terminus. We named this new design ‘HaloCaMP’ (Fig. 1d, Supplementary Fig. 2a,b). These initial prototypes were subjected to mutagenesis targeting the linkers between cpHaloTag and the peptide (L1) and CaM (L2), which led to two calcium sensors: HaloCaMP1a, containing the MLCK peptide, and HaloCaMP1b with the CKK peptide. To explore the modularity of this chemigenetic scaffold, we followed two distinct strategies to produce chemigenetic voltage indicators were also successful. We first inserted cpHaloTag into the loop connecting the third and fourth alpha helices of a VSD (Fig. 1h-i, Supplementary Fig. 2c). We called this sensor scaffold HASAP by analogy with the GFP-based sensor ASAP^5^. We also explored a topology with HaloTag directly fused to the C-terminus of a VSD, analogous to the GFP-based sensor Arclight^18^, which we named HArclight (Fig. 1j-k, Supplementary Fig. 2d).

We focused our biophysical characterization efforts on Ca^2+^ sensors (Fig. 1d-g, Fig. 2a-j), which are soluble proteins amenable to *in vitro* experiments. HaloCaMP1a and HaloCaMP1b showed similar λ_ex_ (640–642 nm) and λ_em_ (653–655 nm) when labeled with **JF**_**635**_**-HTL** (Fig. 1e). The ‘HaloCaMP1a_635_’ showed a change in fluorescence over baseline (ΔF/F_0_) of 5.0 and a high affinity for calcium (*K*_d_ = 190 nM; Fig. 1f). At high [Ca^2+^], HaloCaMP1a_635_ (ε_sat_ = 96,000 M^−1^cm^−1^; Φ_sat_ = 0.78) was 1.7× brighter than GCaMP6s (ε_sat_ = 70,000 M^−1^cm^−1^; Φ_sat_ = 0.64) with fluorescence spectrum red-shifted by 140 nm.^19^ HaloCaMP1b_635_, based on the CKK peptide, showed even higher sensitivity and Ca^2+^ affinity (ΔF/F_0_ = 9.2; *K*_d_ = 43 nM; Fig. 1f) stemming from lower F_0_. To gain insight into the mechanism of fluorescence change, we solved a crystal structure of Ca^2+^-saturated HaloCaMP1b_635_ (Fig. 1g, Supplementary Table 1). The cpHaloTag domain retains the structure of the parent HaloTag with the zwitterionic fluorophore on the surface of the protein. The Ca^2+^-CaM-peptide complex does not sit in immediate proximity to the fluorophore location, however, suggesting that the conformational coupling to fluorescence change results primarily from rearrangement of the linkers.

**Fig. 2.**
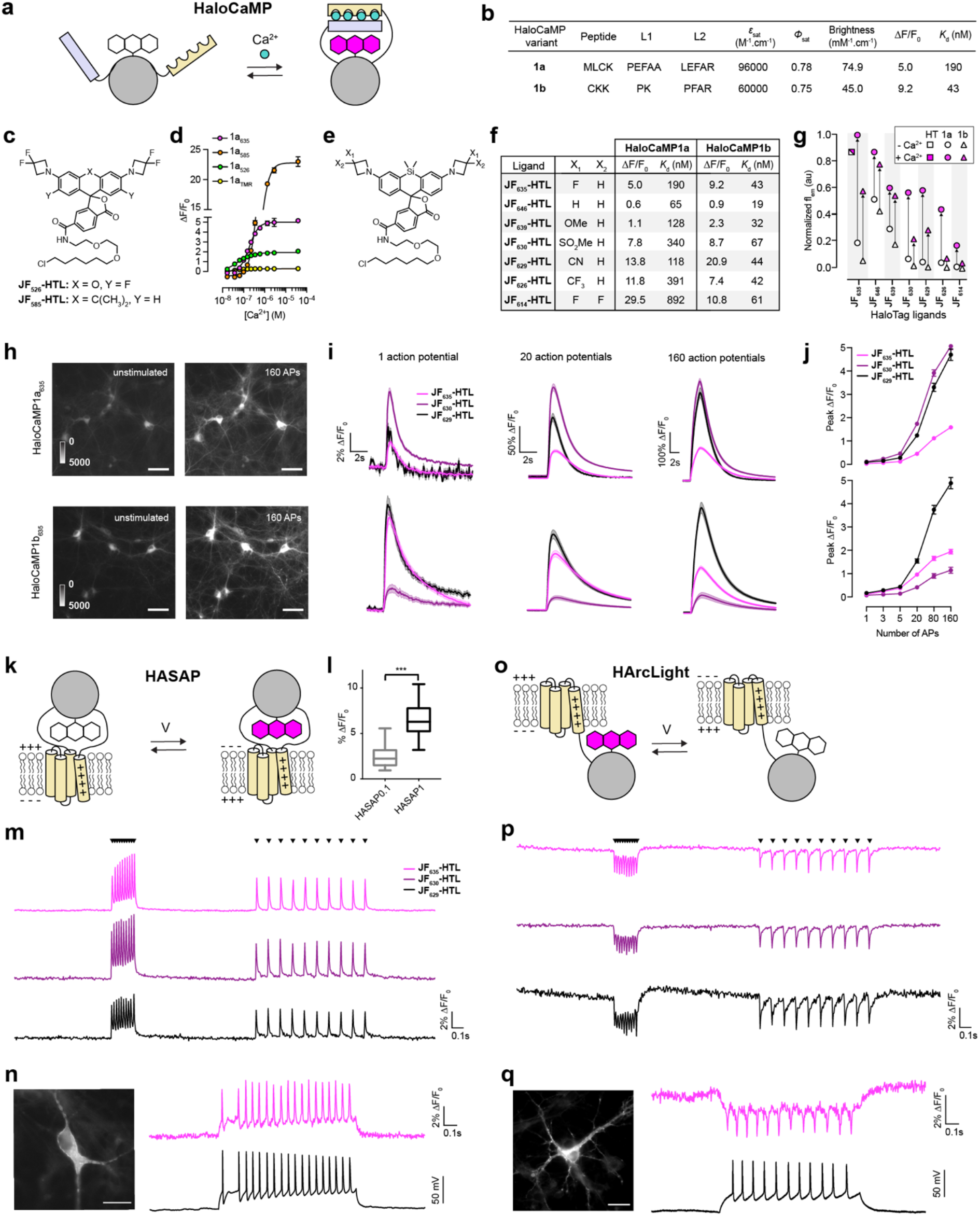
Performance of chemigenetic indicators. **a**, Schematic of HaloCaMP function **b**, Properties of HaloCaMP variants 1a and 1b labeled with **JF**_**635**_**-HTL**. **c**, Structures of **JF**_**526**_**-HTL** and **JF**_**585**_**-HTL**. **d**, Calcium titrations of HaloCaMP1a labeled with different fluorophore ligands. Mean and s.d. for *n* = 2 independent titrations. **e**, General structure of Si-rhodamine HaloTag ligands. **f**, Ca^2+^ binding properties of HaloCaMP1a and 1b bound to Si-rhodamine ligands. **g**, Relative Ca^2+^-free and Ca^2+^-saturated fluorescence of HaloTag (HT), HaloCaMP1a and 1b bound to Si-rhodamine HaloTag ligands; values were normalized to Ca^2+^-saturated HaloCaMP1a_635_. **h**, Cultured rat hippocampal neurons expressing HaloCaMP1a (top panels) or 1b (bottom panels) and labeled with **JF**_**635**_**-HTL**, unstimulated (left panels) and upon 160 APs stimulation (right panels); scale bars: 50 μm. **i**, Fluorescence response of HaloCaMP1a (top) and 1b (bottom) labeled with Si-rhodamine HaloTag ligands to different numbers of action potentials at 80 Hz. Mean and s.e. for *n* = 75 to 120 neurons. **j**, Peak ∆F/F_0_ as a function of the number of APs. Mean and s.e. for *n* = 75 to 120 neurons. **k**, Schematic of HASAP function. **l**, Box plot of fluorescence change of HASAP0.1 (∆F/F_0_ = 2.7 ± 0.4%, *n* = 15 neurons) and HASAP1 (∆F/F_0_ = 6.3 ± 0.4%, *n* = 19 neurons) to field stimulation in neurons. **m**, Fluorescence response of HASAP1 in cultured hippocampal neurons labeled with Si-rhodamine HaloTag ligands to a train of field stimuli eliciting 10 action potentials at 50 Hz followed by 10 action potentials at 10 Hz. Field stimulus timing shown with inverted triangles. **n**, Image of fluorescence in a rat hippocampal neuron in culture (bottom left), and simultaneous fluorescence (magenta) and voltage (black) recording during current injection from a whole-cell patch electrode from a neuron expressing HASAP1 labeled with **JF**_**635**_**-HTL**. Representative of *n* = 4 neurons; scale bar: 20 μm. **o**, Schematic of HArcLight function. **p**, Fluorescence response of HArcLight1 in cultured hippocampal neurons labeled with Si-rhodamine HaloTag ligands to a train of field stimuli eliciting 10 action potentials at 50 Hz followed by 10 action potentials at 10 Hz. Field stimulus timing shown with inverted triangles. **q**, image of fluorescence in a rat hippocampal neuron in culture (bottom left), and simultaneous fluorescence (magenta) and voltage (black) recording during current injection from a whole-cell patch electrode from a neuron expressing HArclight1 labeled with **JF**_**635**_**-HTL**. Representative of *n* = 5 neurons; scale bar: 20 μm.

A useful feature of chemigenetic indicators is the ability to change the small-molecule component to complement protein mutagenesis and allow additional control over sensor properties. We first explored changing spectral properties by testing the yellow and orange fluorogenic HaloTag ligands **JF**_**526**_-HaloTag ligand (λ_ex_/λ_em_ = 526/550 nm)^20^ and **JF**_**585**_-HaloTag ligand (λ_ex_/λ_em_ = 585/609 nm)^15^ as HaloCaMP conjugates (Fig. 2c). Both compounds efficiently labeled HaloCaMP1a and resulted in fluorescence increase with Ca^2+^, with ΔF/F_0_ = 2.0 for the **JF**_**526**_ conjugate and ΔF/F_0_ = 23.2 for the **JF**_**585**_ labeled protein (Fig. 2d). In contrast, we observed no significant Ca^2+^-dependent fluorescence change with the TMR-based ligand (Fig. 2d), consistent with the hypothesis that the HaloCaMP sensor functions by shifting dyes from the lactone to the zwitterion form. Dyes that are already largely shifted to the zwitterionic form, such as TMR, are not significantly modulated by the conformational changes in the protein.

After establishing this flexibility with existing fluorogenic rhodamine dyes, we explored fine-tuning of the small-molecule fluorophores to change sensor properties. We previously reported a general strategy to modulate the *K*_L-Z_ of Janelia Fluor dyes by introducing substituents on the azetidine rings.^15^ We reasoned that minor alterations in the *K*_L-Z_ of Si-rhodamine ligands such as **JF**_**635**_**-HTL** would allow fine-tuning of the sensor properties while retaining far-red excitation and emission. To test this hypothesis, we synthesized a family of novel azetidine-substituted Si-rhodamines and their corresponding HaloTag ligands, using the Pd-catalyzed cross-coupling from a common intermediate (Fig. 2e, Supplementary Fig. 3).^21^ As expected, azetidine substitution elicited only small shifts in λ_ex_ (614–639 nm) and λ_em_ (631–656 nm), with λ_ex_ showing good correlation with the Hammet inductive substituent constants (σ_I_)^22^ for the different azetidine substituents (Supplementary Fig. 4, Supplementary Tables 2,3). The free dyes and unbound HaloTag ligands exhibited low extinction coefficients in aqueous buffer; binding to purified HaloTag protein resulted in a large increase in absorption and fluorescence (Supplementary Fig. 4e).

We then evaluated the performance of these dyes bound to HaloCaMP. All of the new HaloTag ligands efficiently labeled HaloCaMP1a and HaloCaMP1b, and the conjugates were functional sensors, showing fluorescence increase with increasing [Ca^2+^] (Fig. 2f,g, Supplementary Fig. 5). Sensor properties—ΔF/F_0_, *K*_d_ and fluorescence intensity—varied substantially over the series of modified dyes, producing a range of finely tuned indicators. Generally, increasing the electron withdrawing capacity of the azetidine substituents increased ΔF/F_0_, decreasing both F_0_ and F_sat_. **JF**_**646**_ and **JF**_**639**_ provide high Ca^2+^-free fluorescence, a useful property for imaging small or sparse features. In contrast, **JF**_**626**_ and **JF**_**614**_ exhibit low baseline fluorescence and high sensitivity, a desirable feature to achieve low background in densely labeled samples. Ligands **JF**_**635**_, **JF**_**630**_ and **JF**_**629**_ offer a good compromise between brightness and sensitivity.

We tested the ability of our chemigenetic calcium and voltage indicators to detect neuronal activity. HaloCaMP, HASAP, and HArclight could each be expressed in cultured rat hippocampal neurons and successfully labeled with **JF**_**635**_**-HTL** (Fig. 2h-q, Supplementary Fig. 6). To quantify the fluorescence responses, action potentials (AP) were evoked by electrical stimulation.^23^ HaloCaMP1a_635_ and HaloCaMP1b_635_ showed large fluorescence increase upon field electrode stimulation, and could detect single APs with ΔF/F_0_ of 2.3 ± 0.3% and 8.4 ± 0.6% respectively (Fig. 2i,j). Labeling with ligands **JF**_**630**_**-HTL** or **JF**_**629**_**-HTL** also resulted in bright signal and fluorescence increase upon stimulation. Consistent with *in vitro* measurements, **JF**_**629**_**-HTL** resulted in significantly larger ΔF/F_0_ in response to a high-frequency train of action potentials. In contrast, **JF**_**630**_**-HTL** showed higher sensitivity with HaloCaMP1a. The voltage indicators HASAP0.1 and HArclight1 were also capable of clearly detecting action potentials in single imaging trials from neurons (Fig. 1i,k). To further improve the HASAP scaffold, we explored mutagenesis of the inter-domain linkers of HASAP0.1 and identified a R467G mutation at the junction of cpHaloTag and helix S4 of the VSD that more than doubled the voltage sensitivity of the indicator (Fig. 2l). HASAP1 showed fluorescence increases of 4-6% per neuron action potential spike (Fig. 2m), while HArclight1 gave 3-4% decreases per spike (Fig. 2p). Each voltage sensor worked well when loaded with **JF**_**635**_**-HTL, JF**_**630**_**-HTL** or **JF**_**629**_**-HTL** (Fig. 2m,p). Current injection during simultaneous whole-cell patch clamp electrophysiology and fluorescence imaging of HASAP1 and HArclight1 demonstrated that each of these voltage indicators had sufficient sensitivity and response speed to faithfully track membrane potential changes in neurons (Fig. 2n,q).

In conclusion, we establish HaloTag combined with environmentally sensitive rhodamine dyes and sensor protein domains as a new scaffold for fluorescent indicators. Using this scaffold, we successfully engineered bright, far-red calcium and voltage indicators suitable for detecting single action potentials in neurons. Additionally, the diversity of available synthetic fluorophore ligands offers the possibility to tailor the sensor properties (color, sensitivity, brightness, affinity) to specific applications without re-engineering of the protein scaffold. Ultimately, this approach can be extended to other sensing motifs, and provides a blueprint for the design of fluorescent conformational biosensors with the potential to overcome limitations of current FP-based sensors.

## Acknowledgements

We acknowledge the Molecular Biology, Cell Culture and Virus Production facilities at Janelia for assistance. This work was supported by the Howard Hughes Medical Institute. B.M. holds a postdoctoral fellowship from the Research Foundation-Flanders (FWO Vlaanderen).

## Author Contributions

C.D., A.S.A., L.D.L. and E.R.S. conceived the project and wrote the manuscript. C.D. contributed organic synthesis, protein engineering, *in vitro* characterization of proteins, x-ray crystallography, and neuron imaging. A.S.A. contributed protein engineering, neuron imaging, and electrophysiology. H.K.B. contributed protein engineering, in vitro characterization, and neuron imaging. A.B. contributed organic synthesis and x-ray crystallography. N.F. contributed organic synthesis. B.M. and M.C. contributed protein engineering and neuron imaging. L.D.L. and E.R.S. directed the project.

## Competing Interests

C.D., A.S.A., H.K.B., L.D.L. and E.R.S. have filed patent applications on chemigenetic indicators and azetidine-substituted fluorophores.

