## Supplementary Information for "Bright and tunable far-red chemigenetic indicators"

#### SUPPLEMENTARY INFORMATION CONTENTS:

Methods – pages S2-S4

Supplementary Figures – pages S5-S13

Supplementary Tables – pages S14-S16

Synthesis and characterization for all new compounds – pages S17-S37

Supplementary References – page S38

#### METHODS

**Reagent availability.** Plasmids have been deposited at Addgene (www.addgene.org) as follows: pAAV-synapsin-HaloCaMP1a (plasmid #138327), pAAV-synapsin-HaloCaMP1b (plasmid #138328), pCAG-HASAP1 (plasmid #138325), pCAG-HArcLight1 (plasmid #138326). Other materials are available upon reasonable request from the authors.

**Chemical synthesis.** Commercial reagents were obtained from reputable suppliers and used as received. All solvents were purchased in septum-sealed bottles under inert atmosphere. Reaction under inert atmosphere were sealed with septa through which an argon atmosphere was introduced. Reactions were conducted in round-bottomed flasks or septum-capped crimp-top vials containing Teflon-coated magnetic stir bars. Heating of reactions was performed with a stirring hotplate equipped with a thermometer to maintain the indicated temperatures.

Reactions were monitored by thin layer chromatography (TLC) on precoated glass plates (silica gel 60 $F_{254}$) or by LC–MS (Phenomenex Kinetex 2.1 mm × 30 mm 2.6 µm C18 column; 5 to 10 µL injection; 5– 98% $MeCN/H_2O$, linear gradient, with constant 0.1% v/v $HCO_2H$ additive; 6 min run; 0.5 mL/min flow; ESI; positive ion mode). TLC chromatograms were visualized by UV illumination. Reaction products were purified by flash chromatography on an automated purification system using pre-packed silica gel columns or by preparative HPLC (Phenomenex Gemini – NX 30 × 150 mm 5 µm C18 column). Analytical HPLC analysis was performed with an Agilent Eclipse XDB 4.6 × 150 mm 5 µm C18 column. High-resolution mass spectra were obtained from the High Resolution Mass Spectrometry Facility at the University of Iowa.

NMR spectra were recorded on a 400 MHz spectrometer. Deuterated solvents were used as purchased. $^{1}H$ and $^{13}C$ chemical shifts (δ ppm) were referenced to the residual solvent peaks,[1] and $^{19}F$ chemical shifts (δ ppm) were referenced to $CFCl_3$ added to the tube as a standard. Data for $^{1}H$ NMR and $^{19}F$ NMR spectra are reported by chemical shift (δ ppm), multiplicity (s = singlet, d = doublet, t = triplet, q = quartet, p = pentuplet, dd = doublet of doublets, dt = doublet of triplets, m = multiplet, br s = broad signal), coupling constant (Hz), integration. Data for $^{13}C$ NMR spectra are reported by chemical shift (δ ppm) with hydrogen multiplicity (C, CH, $CH_2$, $CH_3$) obtained from DEPT spectra.

**Molecular biology.** Generally, cloning was done by restriction enzyme digest of plasmid backbones, PCR amplification of inserted fragments, and isothermal assembly to combine them, followed by Sanger sequencing to verify DNA sequences. For the preparation of viruses, plasmid DNA was purified by the Janelia Molecular Biology Facility and AAVs were prepared by the Janelia Virus Services.

**Protein expression and purification.** T7express *E. coli* cells transformed with pRSET plasmid containing HaloTag-EGFP, HaloCaMP1a-EGFP, HaloCaMP1b-EGFP, HaloTag, HaloCaMP1a, or HaloCaMP1b were grown in LB medium containing ampicillin for 48 h at 25°C with shaking at 200 rpm. After centrifugation, cells were lysed by sonication and the lysate was clarified by centrifugation. Purification from lysate was achieved by nickel-affinity chromatography, followed by size exclusion chromatography using a Superdex 200 10/300 GL column (GE Healthcare) at a flow rate of 0.5 $ml.min^{-1}$ in 50 mM Tris-HCl, 75 mM NaCl, pH 7.4. Protein concentration was estimated using the extinction coefficient of EGFP ($\varepsilon_{488}$ = 55900 $M^{-1}.cm^{-1}$) or using the extinction coefficient at 280nm estimated from the protein sequence.

**UV–Vis and fluorescence spectroscopy.** All measurements were taken at ambient temperature (23 ± 2°C). Fluorescent molecules were prepared as stock solutions in DMSO and diluted such that the DMSO concentration did not exceed 1% v/v. Spectroscopy was performed using 1-cm path length quartz cuvettes (Starna) or 96-well clear bottom plates (Greiner). Absorption measurements were recorded on a Cary Model 100 spectrometer (Varian). Fluorescence measurements spectra were recorded on a Cary Eclipse fluorometer (Varian) or M100 Pro plate reader (Tecan). Data was analyzed and graphs were plotted using Prism (GraphPad). Absolute quantum yields ($\Phi$) were measured using a Quantaurus-QY spectrometer (model C11374) from Hamamatsu. Measurements were carried out using dilute samples (A < 0.1) and self-absorption corrections were performed using the instrument software.[2] For Si-rhodamine dyes, measurements were made in 10 mM HEPES pH = 7.4. For the corresponding HaloTag ligands, 0.1 $mg.mL^{-}$

[1] CHAPS was added to the buffer. For measurements in the presence of HaloTag protein, the dye was incubated with 1.5 eq of purified HaloTag protein (100 µM solution in 75 mM NaCl, 50 mM Tris-HCl, pH 7.4) for 2 h at room temperature.
For measurements with HaloCaMP protein, the dye was incubated with 1.5 eq of purified HaloCaMP protein (100 µM solution in 75 mM NaCl, 50 mM Tris-HCl, pH 7.4) overnight at room temperature; measurements were then made in a commercial EGTA/Ca·EGTA buffer system (Invitrogen) to which 0.1 mg.mL$^{-1}$ CHAPS was added.

**Calcium titrations**. The HaloTag ligands were incubated with 1.5 eq of purified HaloCaMP1a or HaloCaMP1b overnight to guarantee complete binding of the dye-ligand. Calcium titrations were performed in a commercial EGTA/ Ca·EGTA buffer system (Invitrogen) following the associated protocol. Briefly, different proportions of EGTA buffer (30 mM MOPS pH 7.2, 10 mM EGTA, 100 mM KCl) or Ca·EGTA buffer (30 mM MOPS pH 7.2, 10 mM Ca·EGTA, 100 mM KCl) were mixed to give solutions with different free [$Ca^{2+}$]. Fluorescence emission was measured on a plate reader. For each fluorophore, $\lambda_{ex}$ and $\lambda_{em}$ were adjusted to the values determined for the HaloTag-bound fluorophore. All the calcium titrations were performed in duplicate.

**Imaging HaloCaMP in primary neuron culture**. Primary rat hippocampal neurons were prepared as described previously and infected with AAV viruses. Stock solutions of the different Janelia Fluor ligands were prepared at C = 1 mM in DMSO. Cultured neurons were incubated with the cell-permeant dyes at 37°C for 30 min at a final concentration C = 1 µM before washing twice with imaging buffer containing 145 mM NaCl, 2.5 mM KCl, 10 mM glucose, 10 mM HEPES, pH 7.4, 2 mM $CaCl_2$ and 1 mM $MgCl_2$. Synaptic blockers (10 µM CNQX, 10 µM CPP, 10 µM GABAZINE, and 1 mM MCPG) were added prior to imaging.
Wide-field imaging was performed on an inverted Nikon Eclipse Ti2 microscope equipped with a SPECTRA X light engine (Lumencore) with a 20X objective (NA = 0.75, Nikon), and imaged onto a sCMOS camera (Hamamatsu ORCA-Flash 4.0). A FITC filter set (475/50 nm (excitation), 540/50 nm (emission), 506LP dichroic mirror (FITC-5050A-000; Semrock)) was used to image GFP. A quad bandpass filter (set number: 89000, Chroma) with 645/30 nm (excitation), 705/72 nm (emission), and 660nm dichroic mirror (89100bs; Chroma) was used to image all Si-rhodamine derivatives. Action potentials (APs) were evoked by field stimulation with a custom-built platinum wire electrode inserted in the medium, controlled by a high current isolator (A385, World Precision Instruments) set at 90 mA. We stimulated trains of APs from 1 to 160 APs and acquired time-lapse images before, during and after stimulation. Images were processed in ImageJ/Fiji. $Ca^{2+}$-dependent fluorescence change was measured for single hand-segmented neurons, on 2 to 4 well replicates and multiple fields of view per well.

**Field stimulation of HASAP and HArclight in primary neuron culture.** A stimulus isolator (A385, World Precision Instruments) with platinum wires was used to deliver field stimuli (50V, 1 ms) to elicit or action potentials in cultured neurons as described previously[3]. The stimulation was controlled using Wavesurfer and timing was synchronized with fluorescence acquisition using Wavesurfer and a National Instruments PCIe-6353 board. To measure change of HASAP or HArcLight fluorescence response to action potentials over time, neurons expressing these sensors were labeled with the different JF HaloTag ligands and imaged with a 40x objective at 400 Hz using the same imaging setup as described for HaloCaMP. We applied field electrode stimulations to induce a train of 10 single action potentials at 50Hz, followed by a train of 10 action potentials at 10 Hz. Images were processed in ImageJ. Voltage-dependent fluorescence changes were measured for hand-segmented neurons, on 3 well replicates and multiple fields of view per well.

**Electrophysiology of HASAP and HArclight in primary neuron culture.** Filamented glass micropipettes (Sutter Instruments) were pulled to a tip resistance of 4 – 6 MΩ. Pipettes were positioned with a MPC200 manipulator (Sutter Instruments). Whole cell voltage clamp and current clamp recordings were acquired using an EPC800 amplifier (HEKA), filtered at 10 kHz with the internal Bessel filter, and digitized using a National Instruments PCIe-6353 acquisition board at 20 kHz. Data were acquired from cells with access resistance < 25 MΩ. WaveSurfer software was used to generate the various analog and digital waveforms to control the amplifier, camera, light source, and record voltage and current traces. All electrophysiology measurements were performed in imaging buffer: 145 NaCl, 2.5 KCl, 10 glucose, 10 HEPES, pH 7.4, 2 $CaCl_2$, 1 $MgCl_2$, adjusted to 310 mOsm with sucrose. Internal solution for current clamp recordings contained the following (in mM): 130 potassium methanesulfonate, 10 HEPES, 5 NaCl, 1 $MgCl_2$, 1 Mg-ATP, 0.4 Na-GTP, 14 Tris-phosphocreatine, adjusted to pH 7.3 with KOH, and adjusted to 300 mOsm with sucrose. To generate action potentials, current was injected (20 – 200 pA for 1-2 s) and voltage was monitored.

**Crystallography.** For crystallization, purified HaloTag or HaloCaMP1b was incubated with 3 equivalents of TMR-HTL or **$JF_{635}$-HTL** for at least 5 hours and subsequently purified by size exclusion chromatography in 75 mM NaCl, 50 mM Tris-HCl, pH 7.4 to remove unbound dye-ligand. Crystallization was carried out at ambient temperature (23 ± 2°C) and crystals were grown using the hanging-drop vapor diffusion method in 24-well plates. HaloTag-TMR (10 $mg.mL^{-1}$) was crystallized by mixing with a precipitant solution of 0.2M $MgCl_2$, 0.1M Tris pH 8.5, 20% PEG 8000. $HaloCaMP1b_{635}$ (12 $mg.mL^{-1}$) crystallized 10 days after mixing with a precipitant solution consisting of 30% (v/v) PEG 200, 100 mM MES/ Sodium hydroxide pH 6.0 and 5% (w/v) PEG 3000 using 4 µL protein solution to 1.5 µL precipitant drop ratio. Magenta-colored crystals of HaloTag-TMR were cryoprotected by a quick soak in precipitant solution containing 30% glycerol. Blue $HaloCaMP1b_{635}$ crystals were plunged into liquid nitrogen for storage and transport. X-ray diffraction data for HaloTag-TMR and $HaloCaMP1b_{635}$ were collected at beamlines 24-ID-C at the Advanced Photon Source and 8.2.1 at the Advanced Light Source, respectively, at 100 K under a cold nitrogen stream. Diffraction data were integrated using iMosflm[4] and scaled using Scala from within the CCP4 suite[5]. Structures were solved by molecular replacement using Phaser[6] with prior structures of HaloTag and calmodulin complexes as search models. Iterative cycles of refinement in Refmac[7] and model rebuilding/adjustment in Coot[8] led to the models described in Supplementary Table 1.

#### SUPPLEMENTARY FIGURES

**Figure S1.** Exploration of circular permutation sites within HaloTag for sensor design. (a) Crystal structure of HaloTag (grey cartoon ribbons) bound to TMR-HaloTag ligand (sticks), illustrating circular permutation sites tested as green spheres labeled with the amino acid number. (b) Schematic of the sensor context for testing HaloTag circular permutations, with cpHaloTag inserted between helices 3 and 4 of a VSD (top), the field stimulus pattern for stimulation of neurons expressing voltage sensors with cpHaloTag variants (middle), and representative fluorescence traces of each voltage sensor variant in stimulated neurons (bottom).

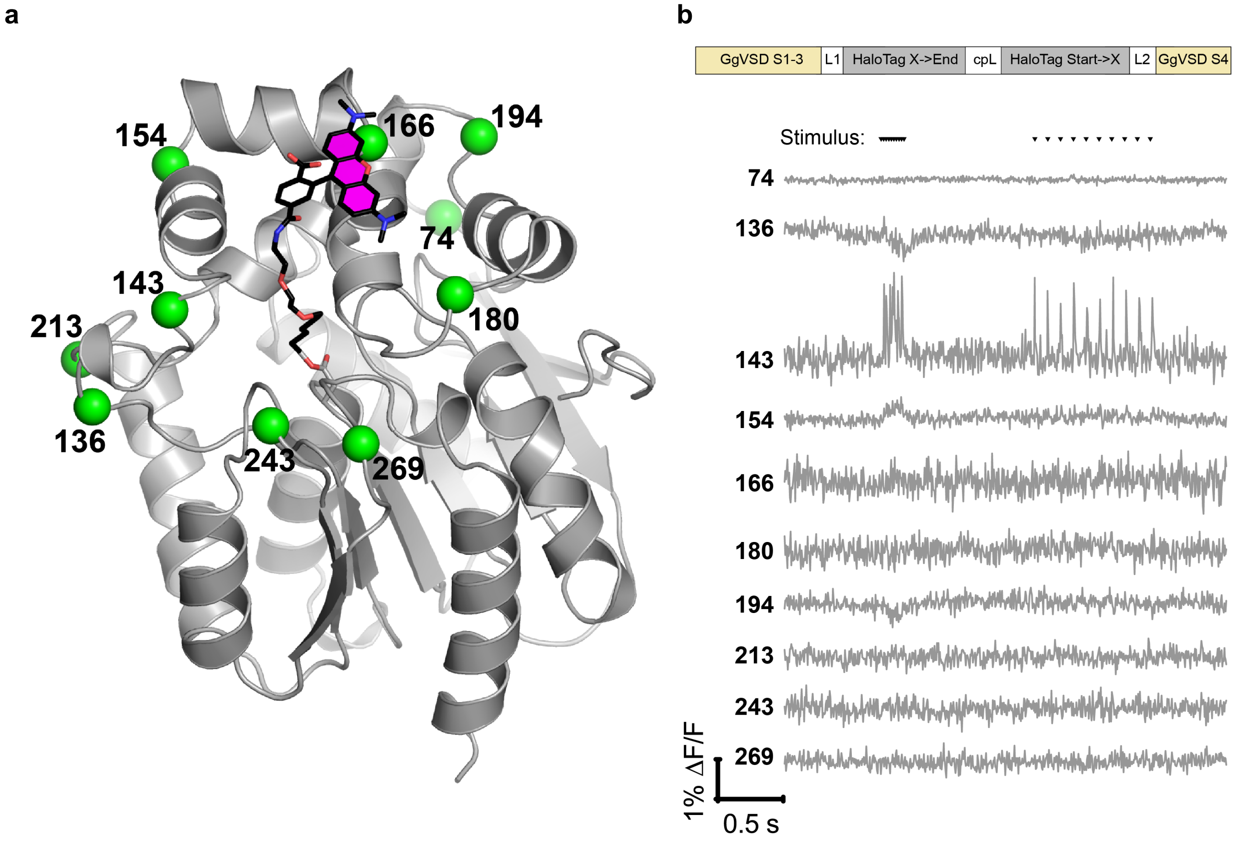

**Figure S2**. DNA and amino acid sequences of HaloCaMP1a (a), HaloCaMP1b (b), HASAP1 (c), and HArcLight1 (d) annotated with sequence features.

#### a – HaloCaMP1a

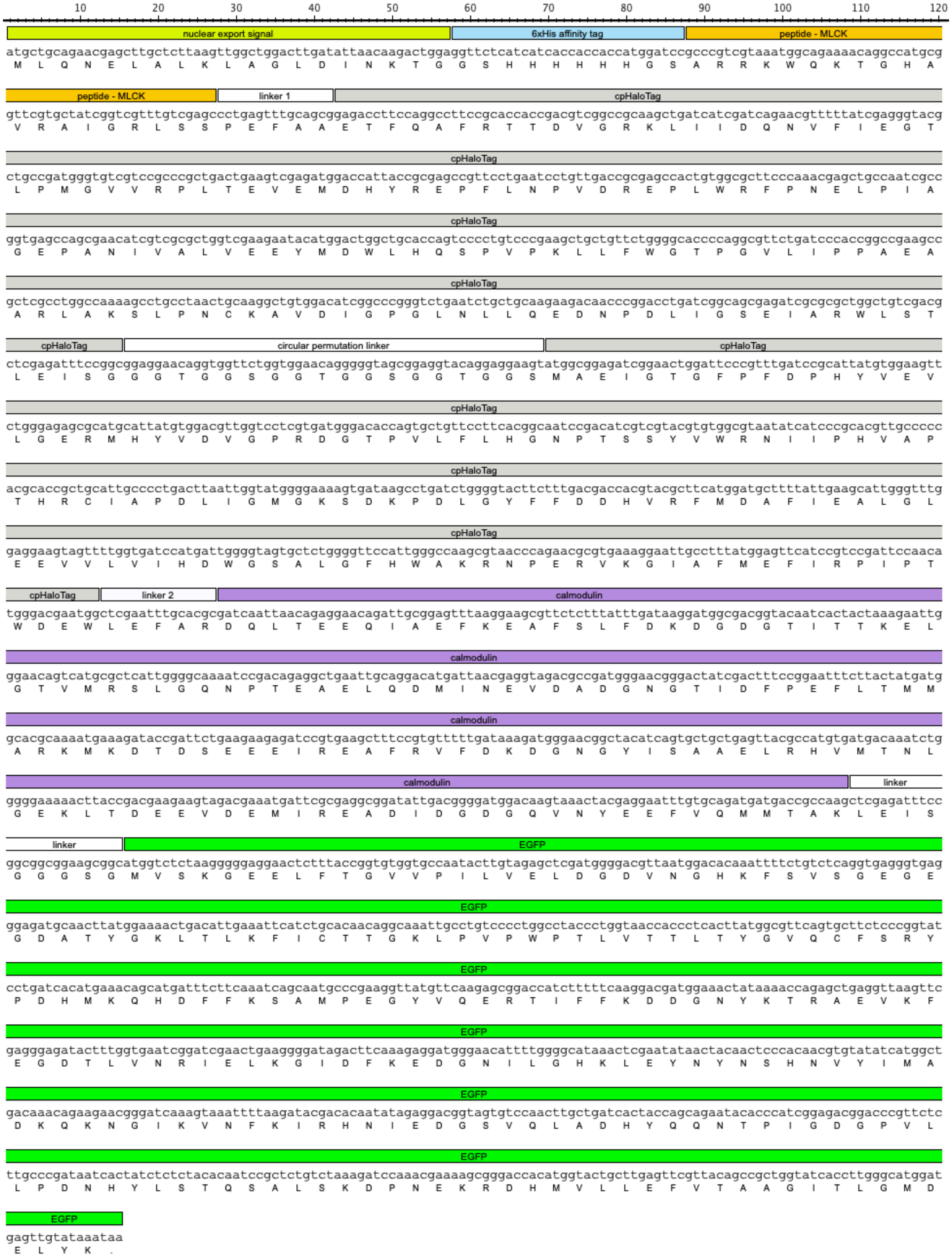

#### b – HaloCaMP1b

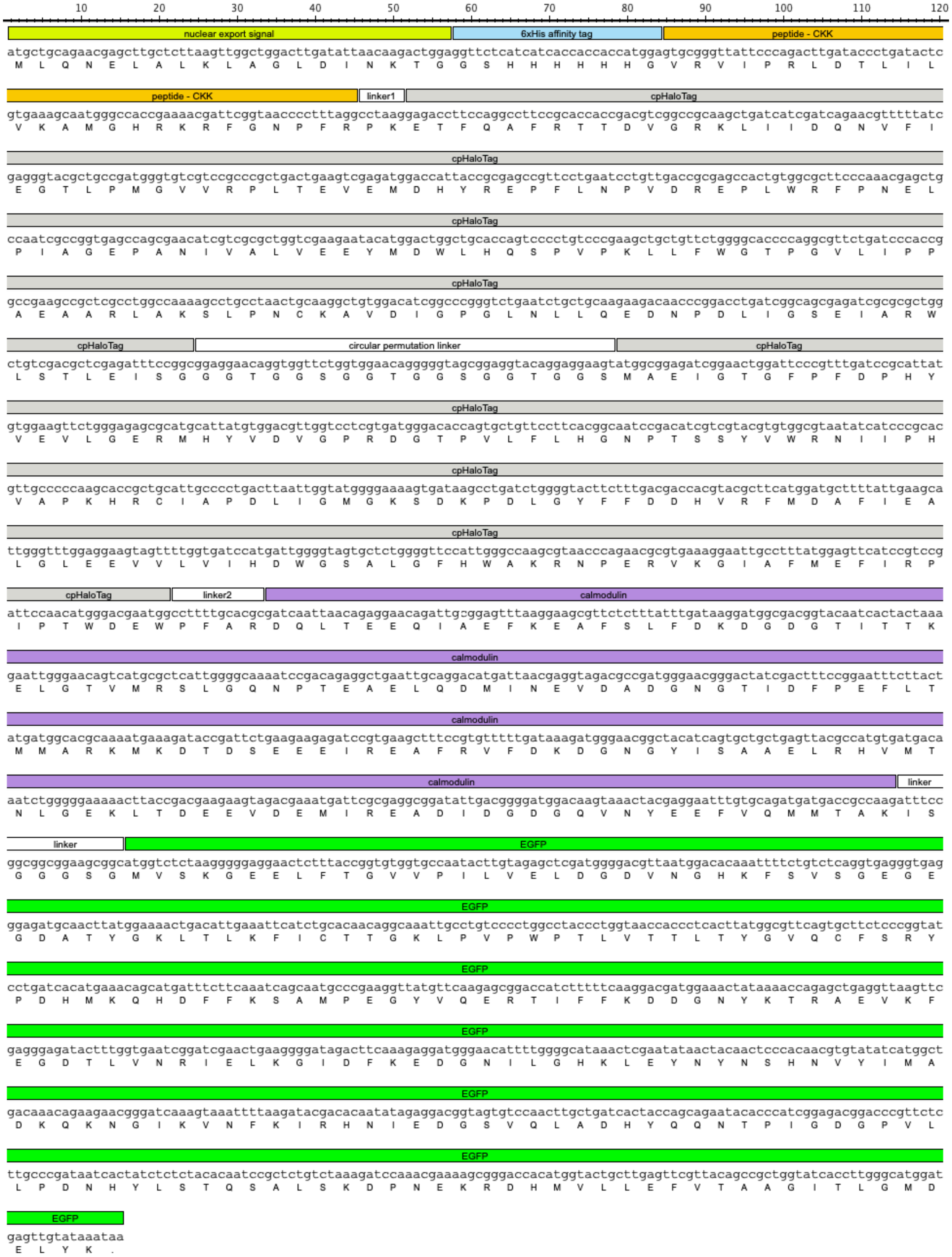

#### c – HASAP1

```
         10        20        30        40        50        60        70        80        90        100       110       120
GgVSD S1-3
atggagacgactgtgaggtatgaacaggggtcagagctcactaaaacttcgagctctccaacagcagatgagcccacgataaagattgatgatggtcgtgatgagggtaatgaacaagac 120
 M  E  T  T  V  R  Y  E  Q  G  S  E  L  T  K  T  S  S  S  P  T  A  D  E  P  T  I  K  I  D  D  G  R  D  E  G  N  E  Q  D

GgVSD S1-3
agctgttccaataccattaggagaaaaatttccccgtttgtgatgtcatttggattcagagtatttggagttgtgcttatcattgtagacatcatagtggtgattgtggatctggccatc 240
 S  C  S  N  T  I  R  R  K  I  S  P  F  V  M  S  F  G  F  R  V  F  G  V  V  L  I  I  V  D  I  I  V  V  I  V  D  L  A  I

GgVSD S1-3
agtgagaagaaaagaggcattagagagattcttgaaggtgtttccctggctatagcactcttcttccttgttgatgttctcatgagagtgtttgttgaaggcttcaagaactatttccgg 360
 S  E  K  K  R  G  I  R  E  I  L  E  G  V  S  L  A  I  A  L  F  F  L  V  D  V  L  M  R  V  F  V  E  G  F  K  N  Y  F  R

GgVSD S1-3                                                                           cpHaloTag
tccaaactgaatactttggatgcagtcatagtagtgggcactctgctaattaatatgacctactccttctctgaccttgctgcctttgcccgcgagaccttccaggccttccgcaccacc 480
 S  K  L  N  T  L  D  A  V  I  V  V  G  T  L  L  I  N  M  T  Y  S  F  S  D  L  A  A  F  A  R  E  T  F  Q  A  F  R  T  T

cpHaloTag
gacgtcggccgcaagctgatcatcgatcagaacgtttttatcgagggtacgctgccgatgggtgtcgtccgcccgctgactgaagtcgagatggaccattaccgcgagccgttcctgaat 600
 D  V  G  R  K  L  I  I  D  Q  N  V  F  I  E  G  T  L  P  M  G  V  V  R  P  L  T  E  V  E  M  D  H  Y  R  E  P  F  L  N

cpHaloTag
cctgttgaccgcgagccactgtggcgcttcccaaacgagctgccaatcgccggtgagccagcgaacatcgtcgcgctggtcgaagaatacatggactggctgcaccagtcccctgtcccg 720
 P  V  D  R  E  P  L  W  R  F  P  N  E  L  P  I  A  G  E  P  A  N  I  V  A  L  V  E  E  Y  M  D  W  L  H  Q  S  P  V  P

cpHaloTag
aagctgctgttctggggcaccccaggcgttctgatcccaccggccgaagccgctcgcctggccaaaagcctgcctaactgcaaggctgtggacatcggcccgggtctgaatctgctgcaa 840
 K  L  L  F  W  G  T  P  G  V  L  I  P  P  A  E  A  A  R  L  A  K  S  L  P  N  C  K  A  V  D  I  G  P  G  L  N  L  L  Q

cpHaloTag                                                                    circular permutation linker
gaagacaacccggacctgatcggcagcgagatcgcgcgctggctgtcgacgctcgagatttccggcgagccaaccactggaggcagcggaggcacaggaggcagcggaggcacaggaggc 960
 E  D  N  P  D  L  I  G  S  E  I  A  R  W  L  S  T  L  E  I  S  G  E  P  T  T  G  G  S  G  G  T  G  G  S  G  G  T  G  G

ci...r cpHaloTag
agcatggcagaaatcggtactggctttccattcgacccccattatgtggaagtcctgggcgagcgcatgcactacgtcgatgttggtccgcgcgatggcacccctgtgctgttcctgcac 1080
 S  M  A  E  I  G  T  G  F  P  F  D  P  H  Y  V  E  V  L  G  E  R  M  H  Y  V  D  V  G  P  R  D  G  T  P  V  L  F  L  H

cpHaloTag
ggtaacccgacctcctcctacgtgtggcgcaacatcatcccgcatgttgcaccgacccatcgctgcattgctccagacctgatcggtatgggcaaatccgacaaaccagacctgggttat 1200
 G  N  P  T  S  S  Y  V  W  R  N  I  I  P  H  V  A  P  T  H  R  C  I  A  P  D  L  I  G  M  G  K  S  D  K  P  D  L  G  Y

cpHaloTag
ttcttcgacgaccacgtccgcttcatggatgccttcatcgaagccctgggtctggaagaggtcgtcctggtcattcacgactggggctccgctctgggtttccactggccaagcgcaat 1320
 F  F  D  D  H  V  R  F  M  D  A  F  I  E  A  L  G  L  E  E  V  V  L  V  I  H  D  W  G  S  A  L  G  F  H  W  A  K  R  N

cpHaloTag                                                                        GgVSD S4
                                                                              R467G
ccagagcgcgtcaaaggtattgcatttatggagttcatccgccctatcccgacctgggacgaatggccagaatttgccgggacagatcagatgcctcaaatggtgacacttttgcgagtt 1440
 P  E  R  V  K  G  I  A  F  M  E  F  I  R  P  I  P  T  W  D  E  W  P  E  F  A  G  T  D  Q  M  P  Q  M  V  T  L  L  R  V

GgVSD S4
ctgcgaatagtgatcctgattcgaatctttcgccttgccagccagaagaaacaactggaggtagtaacataa                                                  1512
 L  R  I  V  I  L  I  R  I  F  R  L  A  S  Q  K  K  Q  L  E  V  V  T  .
```

At amino acid position 467, HASAP0.1=R, HASAP1=G.

#### d – HArcLight1

```
       10        20        30        40        50        60        70        80        90       100       110       120

CiVSD
atggagggattcgacggttcagattttagtcctccagctgatttagttggcgttggcggtgcagtcatgcggaacgtcgttgacgtcacgataaatggtgacgtcactgctccgccgaaa
 M  E  G  F  D  G  S  D  F  S  P  P  A  D  L  V  G  V  G  G  A  V  M  R  N  V  V  D  V  T  I  N  G  D  V  T  A  P  P  K

CiVSD
gcagcgccaagaaaaagtgaatcggtaaagaaagttcattggaatgatgtagaccaaggaccgagtgaaaaaccagagacaagacaggaggaacgaatagatatacccgagatttcaggt
 A  A  P  R  K  S  E  S  V  K  K  V  H  W  N  D  V  D  Q  G  P  S  E  K  P  E  T  R  Q  E  E  R  I  D  I  P  E  I  S  G

CiVSD
ctatggtggggcgagaatgaacatggagtgggcggtgggagaatggagatacctactactggtgtaggtcgcgtccagtttcgtgtccgagcagtgattgatcatctagggatgcgagcc
 L  W  W  G  E  N  E  H  G  V  G  G  G  R  M  E  I  P  T  T  G  V  G  R  V  Q  F  R  V  R  A  V  I  D  H  L  G  M  R  A

CiVSD
tttggagtcttcctaattctcttggacatcatcctcatgatcattgatctcagtcttccaggaaaaagtgaatcttcacaatcctttttatgacgggttggctttggctctttcttgttat
 F  G  V  F  L  I  L  L  D  I  I  L  M  I  I  D  L  S  L  P  G  K  S  E  S  S  Q  S  F  Y  D  G  L  A  L  A  L  S  C  Y

CiVSD
ttcatgctggatttaggattaaggatatttgcctacgggcccaagaatttcttcaccaacccctgggaggttgctgatggcttgattatcgttgtcacattcgtcgtcacgatattttac
 F  M  L  D  L  G  L  R  I  F  A  Y  G  P  K  N  F  F  T  N  P  W  E  V  A  D  G  L  I  I  V  V  T  F  V  V  T  I  F  Y

CiVSD                                                                                                                 H...
actgtgttagatgaatactttcaagaaacaggagccgatggtttggggcagttggttgtgttggcccgtttgctgcgtgtggttcgattagcaagaatattttattcccatcaacaaatc
 T  V  L  D  E  Y  F  Q  E  T  G  A  D  G  L  G  Q  L  V  V  L  A  R  L  L  R  V  V  R  L  A  R  I  F  Y  S  H  Q  Q  I

HaloTag
ggtactggctttccattcgaccccattatgtggaagtcctgggcgagcgcatgcactacgtcgatgttggtccgcgcgatggcacccctgtgctgttcctgcacggtaacccgacctcc
 G  T  G  F  P  F  D  P  H  Y  V  E  V  L  G  E  R  M  H  Y  V  D  V  G  P  R  D  G  T  P  V  L  F  L  H  G  N  P  T  S

HaloTag
tcctacgtgtggcgcaacatcatcccgcatgttgcaccgacccatcgctgcattgctccagacctgatcggtatgggcaaatccgacaaaccagacctgggttatttcttcgacgaccac
 S  Y  V  W  R  N  I  I  P  H  V  A  P  T  H  R  C  I  A  P  D  L  I  G  M  G  K  S  D  K  P  D  L  G  Y  F  F  D  D  H

HaloTag
gtccgcttcatggatgccttcatcgaagccctgggtctggaagaggtcgtcctggtcattcacgactggggctccgctctgggtttccactgggccaagcgcaatccagagcgcgtcaaa
 V  R  F  M  D  A  F  I  E  A  L  G  L  E  E  V  V  L  V  I  H  D  W  G  S  A  L  G  F  H  W  A  K  R  N  P  E  R  V  K

HaloTag
ggtattgcatttatggagttcatccgccctatcccgacctgggacgaatggccagaatttgcccgcgagaccttccaggccttccgcaccaccgacgtcggccgcaagctgatcatcgat
 G  I  A  F  M  E  F  I  R  P  I  P  T  W  D  E  W  P  E  F  A  R  E  T  F  Q  A  F  R  T  T  D  V  G  R  K  L  I  I  D

HaloTag
cagaacgtttttatcgagggtacgctgccgatgggtgtcgtccgcccgctgactgaagtcgagatggaccattaccgcgagccgttcctgaatcctgttgaccgcgagccactgtggcgc
 Q  N  V  F  I  E  G  T  L  P  M  G  V  V  R  P  L  T  E  V  E  M  D  H  Y  R  E  P  F  L  N  P  V  D  R  E  P  L  W  R

HaloTag
ttcccaaacgagctgccaatcgccggtgagccagcgaacatcgtcgcgctggtcgaagaatacatggactggctgcaccagtcccctgtcccgaagctgctgttctggggcaccccaggc
 F  P  N  E  L  P  I  A  G  E  P  A  N  I  V  A  L  V  E  E  Y  M  D  W  L  H  Q  S  P  V  P  K  L  L  F  W  G  T  P  G

HaloTag
gttctgatcccaccggccgaagccgctcgcctggccaaaagcctgcctaactgcaaggctgtggacatcggcccgggtctgaatctgctgcaagaagacaacccggacctgatcggcagc
 V  L  I  P  P  A  E  A  A  R  L  A  K  S  L  P  N  C  K  A  V  D  I  G  P  G  L  N  L  L  Q  E  D  N  P  D  L  I  G  S

HaloTag
gagatcgcgcgctggctgtcgacgctcgagatttccggcgagccaaccacttaa
 E  I  A  R  W  L  S  T  L  E  I  S  G  E  P  T  T  .
```

**Figure S3**. Synthesis of azetidine-substituted Si-rhodamine derivatives (a) and corresponding HaloTag ligands (b).

**a**

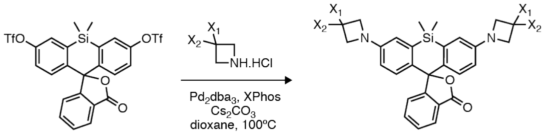


silafluorescein ditriflate

**$JF_{639}$:** $X_1$ = OMe, $X_2$ = H *78%*
**$JF_{630}$:** $X_1$ = $SO_2Me$, $X_2$ = H *80%*
**$JF_{629}$:** $X_1$ = CN, $X_2$ = H *42%*
**$JF_{626}$:** $X_1$ = $CF_3$, $X_2$ = H *76%*
**$JF_{614}$:** $X_1$ = F, $X_2$ = F *24%*

**b**

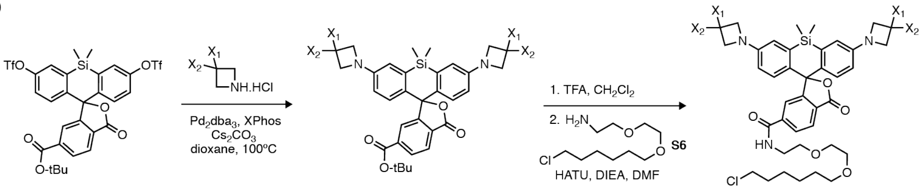


6-*tert*-butoxycarbonyl silafluorescein ditriflate

**S1:** $X_1$ = OMe, $X_2$ = H *94%*
**S2:** $X_1$ = $SO_2Me$, $X_2$ = H *87%*
**S3:** $X_1$ = CN, $X_2$ = H *88%*
**S4:** $X_1$ = $CF_3$, $X_2$ = H *54%*
**S5:** $X_1$ = F, $X_2$ = F *71%*

**$JF_{639}$-HTL:** $X_1$ = OMe, $X_2$ = H *60%*
**$JF_{630}$-HTL:** $X_1$ = $SO_2Me$, $X_2$ = H *65%*
**$JF_{629}$-HTL:** $X_1$ = CN, $X_2$ = H *73%*
**$JF_{626}$-HTL:** $X_1$ = $CF_3$, $X_2$ = H *83%*
**$JF_{614}$-HTL:** $X_1$ = F, $X_2$ = F *75%*

**Figure S4**. (a) Absorption, (b) normalized excitation and (c) normalized fluorescence emission spectra of novel azetidine-substituted Si-rhodamines. (d) Correlation of experimental $\lambda_{ex}$ versus inductive Hammett constants ($\sigma_I$) for Si-rhodamines. (e) Normalized absorption (bold line) and fluorescence (dashed line) spectra of Si-rhodamine HaloTag ligands, in the presence (colored line) or absence (black line) of HaloTag protein. Values were normalized to the HaloTag-bound spectra. All spectra were measured at C = 5 µM in 10 mM HEPES pH = 7.4. In the case of the HaloTag ligands, 0.1 mg.mL$^{-1}$ CHAPS was added to the buffer.

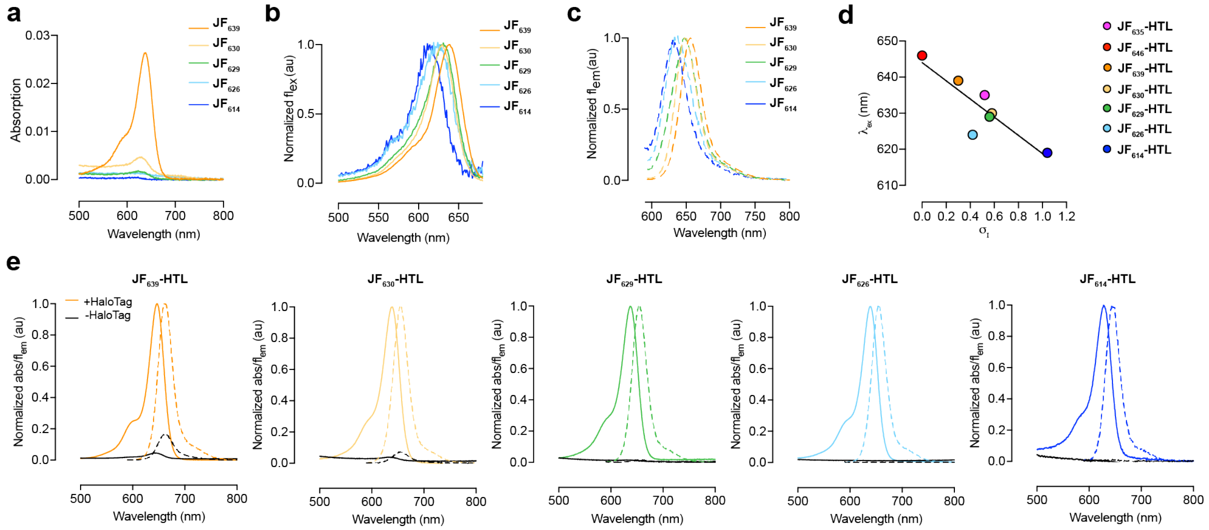

**Figure S5.** Calcium titration curves for HaloCaMP1a (a) and HaloCaMP1b (b) labeled with Si-rhodamine ligands. Mean and s.d. for $n$ = 2 independent titrations.

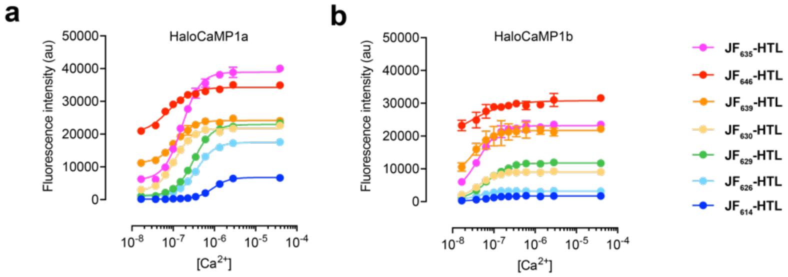

**Figure S6**. Cultured hippocampal neurons expressing HaloCaMP1a-GFP (a) or HaloCaMP1b-GFP labeled with with **$JF_{635}$-HTL** (1 μM, 30 min). GFP channel (left panel), $JF_{635}$ channel (middle panel) and merge (right panel); scale bars: 50 μm.

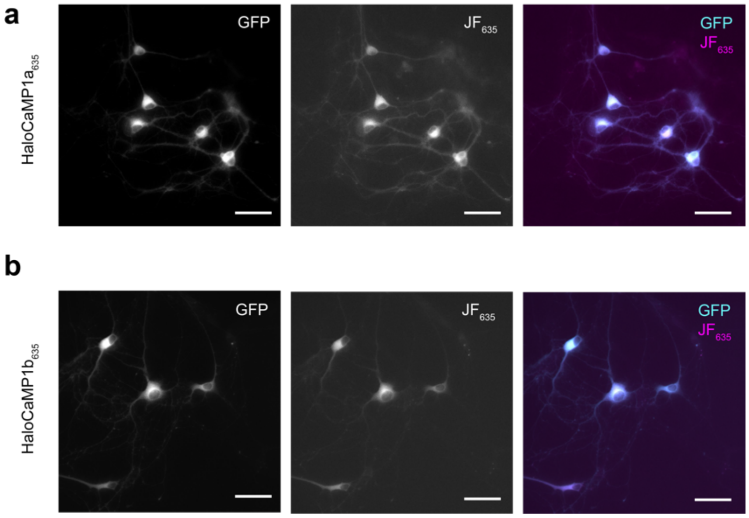

#### SUPPLEMENTARY TABLES

**Table S1.** X-ray diffraction data collection and model refinement statistics.

| | **HaloTag-TMR (PDB 6U32)** | **$Ca^{2+}$-HaloCaMP1b-$JF_{635}$ (PDB 6U2M)** |
|---|---|---|
| **Data collection** | | |
| Space group | $P4_32_12$ | P2 |
| Cell dimensions | | |
| a (Å) | 62.53 | 92.56 |
| b (Å) | 62.53 | 60.66 |
| c (Å) | 164.17 | 122.60 |
| β (degrees) | 90.0 | 91.0 |
| Resolution range (Å)[a] | 62.53 – 1.80 (1.90 – 1.80) | 92.54 – 2.00 (2.11-2.00) |
| Total reflections[a] | 174,075 (25,557) | 421,740 (57,814) |
| Unique reflections[a] | 30,022 (4319) | 90,798 (13,074) |
| Completeness (%)[a] | 97.0 (97.6) | 98.6 (97.6) |
| Redundancy[a] | 5.8 (5.9) | 4.6 (4.4) |
| I/σ[a] | 11.2 (2.4) | 6.5 (1.1) |
| $R_{sym}$ (%)[a,b] | 10.3 (54.5) | 9.6 (69.6) |
| $CC_{1/2}$[a] | 99.8 (69.3) | 98.8 (42.6) |
| **Refinement** | | |
| $R_{work}$ / $R_{free}$ (%)[c] | 15.7/19.3 | 18.8/22.6 |
| Resolution range (Å) | 58.43 – 1.80 | 122.58 – 2.00 |
| Number of atoms (B factor) | | |
| protein | 2350 (28.8) | 7420 (53.1) |
| tetramethylrhodamine-HaloTag ligand | 76 (37.9) | 100 (87.9) |
| chloride ions | 1 (20.8) | 2 (38.9) |
| calcium ions | - | 8 (54.0) |
| water | 177 (36.4) | 203 (47.8) |
| RMSD values | | |
| Bond lengths (Å) | 0.030 | 0.026 |
| Bond angles (degrees) | 2.54 | 2.44 |
| Ramachandran (%) | | |
| Favored/Disallowed | 96.6/0 | 94.7/1.4 |
| Molprobity | | |
| Clashscore (percentile) | 1.7 (100) | 3.8 (99) |
| Molprobity score (percentile) | 1.20 (99) | 2.09 (67) |

[a]The number in parentheses is for the highest resolution shell.
[b]$R_{sym} = \Sigma^i_{hkl} |I^i_{(hkl)} - \langle I_{(hkl)} \rangle| / \Sigma_{hkl} \langle I_{(hkl)} \rangle$, where $I^i_{(hkl)}$ is the $i^{th}$ measured diffraction intensity and $\langle I_{(hkl)} \rangle$ is the mean of the intensity for the miller index (*hkl*).
[c]$R_{work} = \Sigma_{hkl} ||F_o(hkl)| - |F_c(hkl)|| / \Sigma_{hkl} |F_o(hkl)|$. $R_{free} = R_{work}$ for 5% of reflections not included in refinement.

**Table S2.** Photophysical properties of azetidine-substituted Si-rhodamines in 10 mM HEPES, pH = 7.4. NM: not measured.

| Dye | $\lambda_{ex}$ (nm) | $\lambda_{em}$ (nm) | $\varepsilon$ ($M^{-1}.cm^{-1}$) | $\Phi$ |
|---|---|---|---|---|
| **$JF_{635}$** | 635 | 652 | ~400 | 0.56 |
| **$JF_{646}$** | 646 | 664 | 5000 | 0.54 |
| **$JF_{639}$** | 639 | 656 | 5000 | 0.62 |
| **$JF_{630}$** | 630 | 649 | ~700 | NM |
| **$JF_{629}$** | 629 | 648 | <200 | NM |
| **$JF_{626}$** | 626 | 638 | <200 | NM |
| **$JF_{614}$** | 614 | 631 | <200 | NM |

**Table S3.** Photophysical properties of Si-rhodamines HaloTag ligands in the presence or absence of HaloTag protein in 10 mM HEPES pH = 7.4 containing 0.1 mg.mL$^{-1}$ CHAPS. NM: not measured.

| **Ligand** | | **$\lambda_{ex}$ (nm)** | **$\lambda_{em}$ (nm)** | **ε (M$^{-1}$.cm$^{-1}$)** | **Φ** |
|---|---|---|---|---|---|
| **$JF_{635}$-HTL**[9] | - HaloTag | 635 | 652 | ~400 | NM |
| | + HaloTag | 640 | 656 | 81000 | 0.75 |
| **$JF_{646}$-HTL**[9] | - HaloTag | 649 | 666 | 6000 | 0.52 |
| | + HaloTag | 652 | 666 | 95000 | 0.64 |
| **$JF_{639}$-HTL** | - HaloTag | 645 | 658 | 5300 | 0.63 |
| | + HaloTag | 647 | 663 | 120000 | 0.71 |
| **$JF_{630}$-HTL** | - HaloTag | 633 | 657 | 1200 | NM |
| | + HaloTag | 639 | 656 | 32000 | 0.70 |
| **$JF_{629}$-HTL** | - HaloTag | 638 | 655 | <200 | NM |
| | + HaloTag | 638 | 656 | 29000 | 0.81 |
| **$JF_{626}$-HTL** | - HaloTag | 634 | 647 | <200 | NM |
| | + HaloTag | 639 | 654 | 57000 | 0.73 |
| **$JF_{614}$-HTL** | - HaloTag | 622 | 640 | <200 | NM |
| | + HaloTag | 628 | 646 | 7000 | 0.74 |

#### SYNTHESIS AND CHARACTERIZATION FOR ALL NEW COMPOUNDS

**Procedure A:** Synthesis of Si-rhodamines by Pd-catalyzed cross-coupling. The following procedure for **1** is representative. A vial was charged with silafluorescein ditriflate[10] (50 mg, 78 μmol), 3-methoxyazetidine hydrochloride (39 mg, 312 μmol, 4 eq), $Pd_2dba_3$ (7.1 mg, 7.8 μmol, 0.1 eq), XPhos (11.2 mg, 23.4 μmol, 0.3 eq), and $Cs_2CO_3$ (204 mg, 625 mmol, 8 eq). The vial was sealed and evacuated/backfilled with nitrogen (3x). Dioxane (2 mL) was added, and the reaction was flushed again with nitrogen (3x). The reaction was then stirred at 100 °C overnight. It was subsequently cooled to room temperature, diluted with MeOH, deposited onto Celite, and concentrated to dryness. The residue was purified as described.

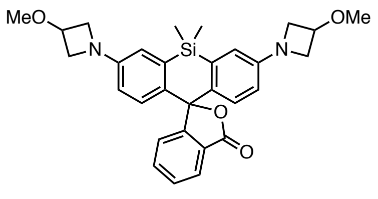


**(1; $JF_{639}$):** Purification by silica gel chromatography (0–35% EtOAc/toluene, linear gradient) afforded **1** (78%) as a light blue solid. $^1$H NMR ($CDCl_3$, 400 MHz) δ 7.96 (d, *J* = 7.6 Hz, 1H), 7.64 (td, *J* = 7.5, 1.2 Hz, 1H), 7.54 (td, *J* = 7.5, 1.0 Hz, 1H), 7.32 – 7.27 (m, 1H), 6.77 (d, *J* = 8.7 Hz, 2H), 6.69 (d, *J* = 2.7 Hz, 2H), 6.28 (dd, *J* = 8.7, 2.7 Hz, 2H), 4.38 – 4.27 (m, 2H), 4.10 (d, *J* = 7.3 Hz, 4H), 3.73 (dt, *J* = 7.7, 4.0 Hz, 4H), 3.32 (s, 6H), 0.61 (s, 3H), 0.58 (s, 3H); $^{13}$C NMR ($CDCl_3$, 101 MHz) δ 170.7 (C), 154.3 (C), 150.4 (C), 137.1 (C), 133.8 (CH), 133.3 (C), 128.9 (CH), 128.1 (CH), 127.0 (C), 125.9 (CH), 124.7 (CH), 116.1 (CH), 112.7 (CH), 91.9 (C), 70.1 ($CH_3$), 58.9 ($CH_2$), 56.2 (CH), 0.5 ($CH_3$), -1.4 ($CH_3$); HRMS (ESI) calcd for $C_{30}H_{33}N_2O_4Si$ $[M+H]^+$ 513.2210, found 513.2202.

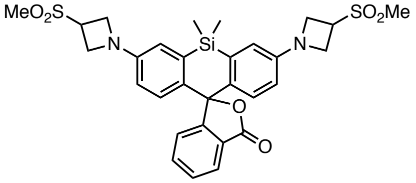


**(2; $JF_{630}$):** Synthesized following procedure A from silafluorescein ditriflate and 3-methylsulfonyl-azetidine hydrochloride. Purification by silica gel chromatography (50–100% EtOAc/hexane, linear gradient) afforded **2** (80%) as a light blue solid. $^1$H NMR ($CDCl_3$, 400 MHz) δ 7.96 (dt, *J* = 7.6, 1.0 Hz, 1H), 7.65 (td, *J* = 7.5, 1.2 Hz, 1H), 7.55 (td, *J* = 7.5, 1.0 Hz, 1H), 7.28 – 7.25 (m, 1H), 6.83 (d, *J* = 8.7 Hz, 2H), 6.70 (d, *J* = 2.7 Hz, 2H), 6.31 (dd, *J* = 8.7, 2.7 Hz, 2H), 4.29 – 4.15 (m, 8H), 4.07 (tt, *J* = 7.5, 5.7 Hz, 2H), 2.96 (s, 6H), 0.62 (s, 3H), 0.59 (s, 3H); $^{13}$C NMR ($CDCl_3$, 101 MHz) δ 170.6 (C), 154.2 (C), 149.2 (C), 137.1 (C), 134.8 (C), 134.1 (CH), 129.1 (CH), 128.2 (CH), 126.6 (C), 126.0 (CH), 124.6 (CH), 116.2 (CH), 113.0 (CH), 91.2 (C), 52.5 ($CH_2$), 52.4 ($CH_2$), 51.7 (CH), 38.3 ($CH_3$), 0.4 ($CH_3$), -1.3 ($CH_3$); HRMS (ESI) calcd for $C_{30}H_{33}N_2O_6SiS_2$ $[M+H]^+$ 609.1549, found 609.1548.

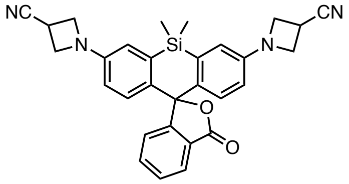


**(3; $JF_{629}$):** Synthesized following procedure A from silafluorescein ditriflate and 3-azetidinecarbonitrile hydrochloride. Purification by HPLC (35–95% $MeCN/H_2O$ + 0.1% TFA additive) afforded **3** (42%) as a light blue solid. $^1$H NMR ($CD_2Cl_2$, 400 MHz) δ 7.93 (dt, *J* = 7.6, 1.0 Hz, 1H), 7.66 (td, *J* = 7.5, 1.2 Hz, 1H), 7.57

(td, *J* = 7.5, 1.0 Hz, 1H), 7.24 (dd, *J* = 7.7, 1.0 Hz, 1H), 6.84 (d, *J* = 8.7 Hz, 2H), 6.72 (d, *J* = 2.7 Hz, 2H), 6.33 (dd, *J* = 8.7, 2.7 Hz, 2H), 4.20 (ddd, *J* = 8.6, 7.1, 1.8 Hz, 4H), 4.13 – 4.03 (m, 4H), 3.60 (tt, *J* = 8.4, 6.1 Hz, 2H), 0.63 (s, 3H), 0.57 (s, 3H); $^{13}$C NMR ($CD_2Cl_2$, 101 MHz) δ $^{13}$C NMR (101 MHz, $CD_2Cl_2$) δ 170.7 (C), 154.8 (C), 150.1 (C), 137.3 (C), 135.0 (C), 134.6 (CH), 129.6 (CH), 128.5 (CH), 126.8 (C), 126.3 (CH), 124.8 (CH), 120.5 (C), 116.6 (CH), 113.4 (CH), 91.4 (C), 55.9 ($CH_2$), 19.1 (CH), 0.4 ($CH_3$), -1.1 ($CH_3$); HRMS (ESI) calcd for $C_{30}H_{27}N_4O_2Si$ $[M+H]^+$ 503.1903, found 503.1899.

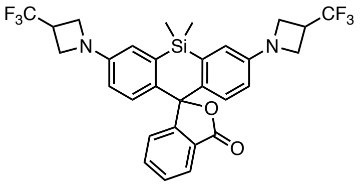


**(4; JF$_{626}$):** Synthesized following procedure A from silafluorescein ditriflate and 3-(trifluoromethyl)azetidine hydrochloride. Purification by silica gel chromatography (0–100% EtOAc/hexane, linear gradient), followed by purification by silica gel chromatography (0–35% EtOAc/toluene) afforded **4** (76%) as a light blue solid. $^1$H NMR ($CDCl_3$, 400 MHz) δ 7.97 (dt, *J* = 7.7, 1.0 Hz, 1H), 7.66 (td, *J* = 7.5, 1.2 Hz, 1H), 7.56 (td, *J* = 7.5, 1.0 Hz, 1H), 7.29 (dd, *J* = 7.7, 0.9 Hz, 1H), 6.82 (d, *J* = 8.7 Hz, 2H), 6.70 (d, *J* = 2.7 Hz, 2H), 6.29 (dd, *J* = 8.7, 2.7 Hz, 2H), 4.14 – 4.04 (m, 4H), 4.01 – 3.91 (m, 4H), 3.49 – 3.29 (m, 2H), 0.62 (s, 3H), 0.60 (s, 3H); $^{19}$F NMR ($CDCl_3$, 376 MHz) = -73.4 (d, $^3J_{HF}$ = 8.8 Hz); $^{13}$C NMR ($CDCl_3$, 101 MHz) δ 170.5 (C), 154.1 (C), 149.5 (C), 137.2 (C), 134.1 (CH), 133.9 (C), 129.1 (CH), 128.2 (CH), 126.5 (q, $^2J_{CF}$ = 81.5Hz, $CF_3$), 126.0 (CH), 124.7 (CH), 115.8 (CH), 112.5 (CH), 91.5 (C), 51.4 ($CH_2$), 51.3 (CH2), 32.8 (q, $^3J_{CF}$ = 32.3 Hz, C), 0.5 ($CH_3$), -1.5 ($CH_3$); HRMS (ESI) calcd for $C_{30}H_{27}N_2O_2SiF_6$ $[M+H]^+$ 589.1746, found 589.1751.

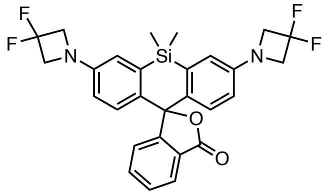


**(5; JF$_{614}$):** Synthesized following procedure A from silafluorescein ditriflate and 3,3-difluoroazetidine hydrochloride. Purification by silica gel chromatography (0–100% EtOAc/hexane, linear gradient), followed by purification by silica gel chromatography (0–35% EtOAc/toluene) afforded **5** (24%) as a light blue solid. $^1$H NMR ($CDCl_3$, 400 MHz) δ 7.98 (dt, *J* = 7.6, 1.0 Hz, 1H), 7.66 (td, *J* = 7.5, 1.2 Hz, 1H), 7.56 (td, *J* = 7.5, 1.0 Hz, 1H), 7.30 – 7.28 (m, 1H), 6.85 (d, *J* = 8.7 Hz, 2H), 6.73 (d, *J* = 2.7 Hz, 2H), 6.34 (dd, *J* = 8.7, 2.8 Hz, 2H), 4.23 (t, $^3J_{HF}$ = 11.8 Hz, 8H), 0.64 (s, 3H), 0.61 (s, 3H); $^{19}$F NMR ($CDCl_3$, 376 MHz) = -99.9 (p, $^3J_{HF}$ = 11.7 Hz) ;$^{13}$C NMR ($CDCl_3$, 101 MHz) δ 170.5 (C), 154.0 (C), 148.7 (t, $^4J_{CF}$ = 2.6 Hz, C), 137.3 (C), 134.8 (C), 134.0 (CH), 129.2 (CH), 128.2 (CH), 126.8 (C), 126.1 (CH), 124.6 (CH), 116.8 (CH), 115.9 (t, $^1J_{CF}$ = 276 Hz, $CF_2$) , 113.6 (CH), 91.2 (C), 63.4 (t, $^2J_{HF}$ = 25.9,$CH_2$), 0.4 ($CH_3$), -1.4 ($CH_3$); HRMS (ESI) calcd for $C_{28}H_{25}N_2O_2SiF_4$ $[M+H]^+$ 525.1621, found 525.1629.

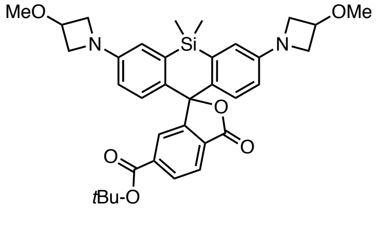


**(S1):** Synthesized following procedure A from 6-*tert*-butoxycarbonylsilafluorescein ditriflate[10] and 3-methoxyazetidine hydrochloride. Purification by silica gel chromatography (0–30% EtOAc/hexane, linear gradient), afforded **S1** (94%) as an off-white solid. $^1$H NMR ($CDCl_3$, 400 MHz) δ δ 8.11 (dd, *J* = 8.1, 1.3 Hz, 1H), 7.96 (d, *J* = 8.0 Hz, 1H), 7.81 (d, *J* = 1.2 Hz, 1H), 6.85 (d, *J* = 8.7 Hz, 2H), 6.68 (d, *J* = 2.7 Hz, 2H),

6.32 (dd, $J$ = 8.7, 2.7 Hz, 2H), 4.38 – 4.27 (m, 2H), 4.16 – 4.04 (m, 4H), 3.78 – 3.68 (m, 4H), 3.32 (s, 6H), 1.55 (s, 9H), 0.65 (s, 3H), 0.58 (s, 3H); $^{13}$C NMR ($CDCl_3$, 101 MHz) δ 170.3 (C), 164.4 (C), 155.4 (C), 150.4 (C), 137.3 (C), 136.2 (C), 132.8 (C), 130.0 (CH), 129.1 (C), 127.7 (CH), 125.7 (CH), 125.1 (CH), 116.1 (CH), 113.1 (CH), 91.7 (C), 82.4 (C), 70.1 ($CH_3$), 58.9 ($CH_2$), 56.2 (CH), 28.2 ($CH_3$), 0.2 ($CH_3$), -0.7 ($CH_3$); HRMS (ESI) calcd for $C_{35}H_{41}N_2O_6Si$ $[M+H]^+$ 613.2734, found 613.2726.

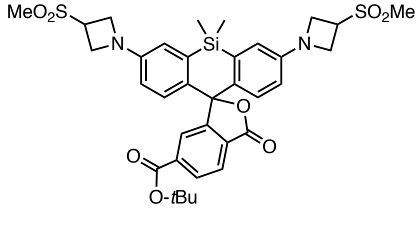


**(S2):** Synthesized following procedure A from 6-*tert*-butoxycarbonylsilafluorescein ditriflate and 3-methylsulfonyl-azetidine hydrochloride. Purification by silica gel chromatography (50–100% EtOAc/hexane, linear gradient), afforded **S1** (87%) as a light blue solid. $^1$H NMR ($CDCl_3$, 400 MHz) δ 8.12 (dd, $J$ = 8.0, 1.3 Hz, 1H), 7.97 (dd, $J$ = 8.0, 0.7 Hz, 1H), 7.79 (t, $J$ = 1.0 Hz, 1H), 6.93 (d, $J$ = 8.7 Hz, 2H), 6.72 (d, $J$ = 2.7 Hz, 2H), 6.38 (dd, $J$ = 8.8, 2.7 Hz, 2H), 4.31 – 4.19 (m, 8H), 4.15 – 4.03 (m, 2H), 2.97 (s, 6H), 1.55 (s, 9H), 0.67 (s, 3H), 0.59 (s, 3H); $^{13}$C NMR ($CDCl_3$, 101 MHz) δ 170.2 (C), 164.3 (C), 155.2 (C), 149.2 (C), 137.5 (C), 136.2 (C), 134.3 (C), 130.2 (CH), 128.7 (C), 127.8 (CH), 125.9 (CH), 124.9 (CH), 116.2 (CH), 113.3 (CH), 90.9 (C), 82.6 (C), 52.5 ($CH_2$), 51.7 (CH), 38.3 ($CH_3$), 28.2 ($CH_3$), 0.1 ($CH_3$), -0.6 ($CH_3$); HRMS (ESI) calcd for $C_{35}H_{41}N_2O_8SiS_2$ $[M+H]^+$ 709.2073, found 709.2074.

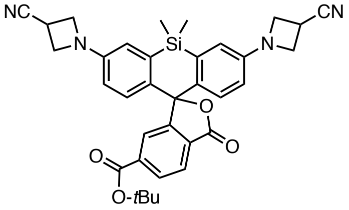


**(S3):** Synthesized following procedure A from 6-*tert*-butoxycarbonylsilafluorescein ditriflate and 3-azetidinecarbonitrile hydrochloride. Purification by silica gel chromatography (0–20% EtOAc/hexane, linear gradient) afforded **S5** (88%) as a light blue solid. $^1$H NMR ($CDCl_3$, 400 MHz) δ 8.13 (dd, $J$ = 8.0, 1.3 Hz, 1H), 7.97 (dd, $J$ = 8.0, 0.7 Hz, 1H), 7.81 (t, $J$ = 1.0 Hz, 1H), 6.91 (d, $J$ = 8.7 Hz, 2H), 6.68 (d, $J$ = 2.7 Hz, 2H), 6.33 (dd, $J$ = 8.7, 2.7 Hz, 2H), 4.25 – 4.17 (m, 4H), 4.14 – 4.02 (m, 4H), 3.59 (tt, $J$ = 8.4, 6.2 Hz, 2H), 1.55 (s, 9H), 0.68 (s, 3H), 0.59 (s, 3H); $^{13}$C NMR ($CDCl_3$, 101 MHz) 170.0 (C), 164.3 (C), 154.9 (C), 149.4 (C), 137.5 (C), 136.3 (C), 134.4 (C), 130.2 (CH), 128.8 (C), 127.8 (CH), 125.9 (CH), 124.9 (CH), 119.7 (C), 116.1 (CH), 113.3 (CH), 90.9 (C), 82.6 (C), 55.3 ($CH_2$), 28.2 ($CH_3$), 18.5 (CH), 0.1 ($CH_3$), -0.7 ($CH_3$); HRMS (ESI) calcd for $C_{35}H_{35}N_4O_4Si$ $[M+H]^+$ 603.2428, found 603.2425.

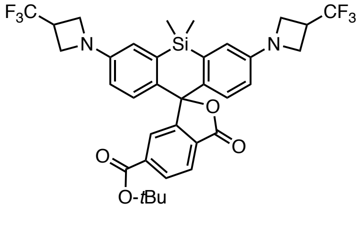


**(S4):** Synthesized following procedure A from 6-*tert*-butoxycarbonylsilafluorescein ditriflate and 3-(trifluoromethyl)azetidine hydrochloride. Purification by silica gel chromatography (0–20% EtOAc/hexane, linear gradient) afforded **S4** (54%) as a light blue solid. $^1$H NMR ($CDCl_3$, 400 MHz) δ 8.12 (dd, $J$ = 8.1, 1.3 Hz, 1H), 7.97 (d, $J$ = 8.0 Hz, 1H), 7.82 (s, 1H), 6.88 (d, $J$ = 8.7 Hz, 2H), 6.68 (d, $J$ = 2.6 Hz, 2H), 6.33 (dd, $J$ = 8.7, 2.7 Hz, 2H), 4.08 (t, $J$ = 8.1 Hz, 4H), 3.98 (dt, $J$ = 7.8, 5.6 Hz, 4H), 3.39 (qt, $J$ = 8.5, 5.8 Hz, 2H),

1.55 (s, 9H), 0.66 (s, 3H), 0.59 (s, 3H); $^{13}C$ NMR ($CDCl_3$, 101 MHz) δ 170.2 (C), 164.4 (C), 155.0 (C), 149.5 (C), 137.4 (C), 136.4 (C), 133.6 (C), 130.1 (CH), 129.0 (C), 127.8 (CH), 125.8 (CH), 125.1 (CH), 125.0 (C), 115.8 (CH), 112.8 (CH), 91.3 (C), 82.5 (C), 51.33 ($CH_2$), 51.30 ($CH_2$), 33.2 (q, $^3J_{CF}$ = 32.1 Hz, C), 28.2 ($CH_3$), 0.2 ($CH_3$), -0.7 ($CH_3$); HRMS (ESI) calcd for $C_{35}H_{35}N_2O_4SiF_6$ $[M+H]^+$ 689.2270, found 689.2282.

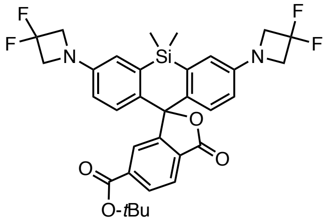


**(S5):** Synthesized following procedure A from 6-*tert*-butoxycarbonylsilafluorescein ditriflate and 3,3-difluoroazetidine hydrochloride. Purification by silica gel chromatography (0–30% EtOAc/hexane, linear gradient), followed by purification by silica gel chromatography (0–20% EtOAc/hexanes) afforded **S5** (71%) as an off-white solid. $^1H$ NMR ($CDCl_3$, 400 MHz) δ 8.13 (dd, $J$ = 8.0, 1.3 Hz, 1H), 7.98 (dd, $J$ = 8.1, 0.8 Hz, 1H), 7.82 (t, $J$ = 1.0 Hz, 1H), 6.93 (d, $J$ = 8.7 Hz, 2H), 6.73 (d, $J$ = 2.7 Hz, 2H), 6.38 (dd, $J$ = 8.7, 2.7 Hz, 2H), 4.24 (t, $J$ = 11.7 Hz, 8H), 1.55 (s, 9H), 0.68 (s, 3H), 0.61 (s, 3H); $^{19}F$ NMR ($CDCl_3$, 376 MHz) = -99.3 (p, $^3J_{HF}$ = 11.9 Hz); $^{13}C$ NMR ($CDCl_3$, 101 MHz) δ 170.0 (C), 164.3 (C), 155.0 (C), 148.7 (t, $^4J_{HF}$ = 2.8 Hz, C), 137.5 (C), 136.5 (C), 134.3 (C), 130.2 (CH), 128.9 (C), 127.9 (CH), 125.9 (CH), 125.0 (C), 116.8 (CH), 115.9 (t, $^1J_{CF}$ = 276 Hz, $CF_2$), 113.9 (CH), 91.0 (C), 82.5 (C), 63.4 (t, $^2J_{HF}$ = 26.0 Hz, $CH_2$), 28.2 ($CH_3$), 0.2 ($CH_3$), -0.7 ($CH_3$); HRMS (ESI) calcd for $C_{33}H_{33}N_2O_4SiF_4$ $[M+H]^+$ 625.2146, found 625.2145.

**Procedure B:** Synthesis of HaloTag ligands. The following procedure for **1-HTL** is representative. **S1** (36 mg, 59 µmol) was taken up in $CH_2Cl_2$ (2 mL) and trifluoroacetic acid (0.25 mL) was added. The reaction was stirred at room temperature overnight. Toluene (3 mL) was added, the reaction mixture was concentrated to dryness and then azeotroped with MeOH three times. The residue was combined with HaloTag($O_2$)amine (**S6**; TFA salt, 30 mg, 89 µmol, 1.5 eq), HATU (34 mg, 89 µmol, 1.5 eq) in DMF (1.5 mL). DIEA (52 µL, 295 µmol, 5.0 eq) was added and the mixture was stirred at room temperature for 4 h. It was subsequently evaporated to dryness and purified as described.

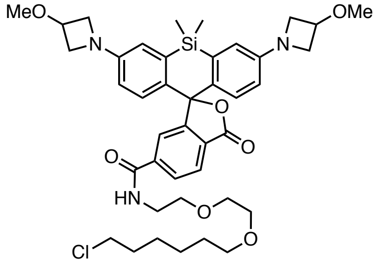


**(1-HTL; JF$_{639}$-HTL):** Purification by silica gel chromatography (30–100% EtOAc/hexanes, linear gradient) provided **1-HTL** (60%) as a light-blue solid. $^1H$ NMR ($CDCl_3$, 400 MHz) δ 7.98 (dd, $J$ = 8.0, 0.7 Hz, 1H), 7.91 (dd, $J$ = 8.0, 1.4 Hz, 1H), 7.68 (t, $J$ = 1.0 Hz, 1H), 6.81 (br s, 1H), 6.76 (d, $J$ = 8.6 Hz, 2H), 6.68 (d, $J$ = 2.7 Hz, 2H), 6.29 (dd, $J$ = 8.7, 2.7 Hz, 2H), 4.37 – 4.29 (m, 2H), 4.13 – 4.07 (m, 4H), 3.76 – 3.70 (m, 4H), 3.66 – 3.60 (m, 6H), 3.56 – 3.52 (m, 2H), 3.50 (t, $J$ = 6.7 Hz, 2H), 3.39 (t, $J$ = 6.7 Hz, 2H), 3.32 (s, 6H), 1.78 – 1.69 (m, 2H), 1.51 (p, $J$ = 6.9 Hz, 2H), 1.44 – 1.35 (m, 2H), 1.34 – 1.23 (m, 2H), 0.64 (s, 3H), 0.57 (s, 3H); Analytical HPLC: $t_R$ = 13.0 min, 99% purity (10–95% MeCN/$H_2O$, linear gradient, with constant 0.1% v/v TFA additive, 20 min run, 1 mL/min flow, detection at 254 nm); HRMS (ESI) calculated for $C_{41}H_{53}ClN_3O_7Si$ $[M+H]^+$ 762.3341, found 762.3352.

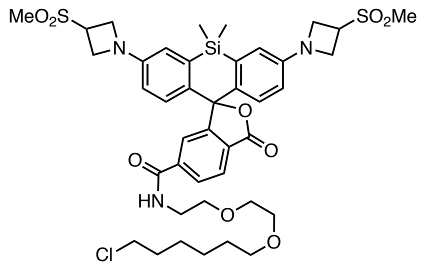


**(2-HTL; $JF_{630}$-HTL):** Synthesized following procedure B from **S2**. Purification by silica gel chromatography (0–4% MeOH/$CH_2Cl_2$, linear gradient), followed by purification by silica gel chromatography (50–100% EtOAc/hexanes, linear gradient) afforded **2-HTL** (65%) as a light blue solid. $^1H$ NMR ($CDCl_3$, 400 MHz) δ.97 (d, *J* = 7.9 Hz, 1H), 7.89 (dd, *J* = 8.0, 1.4 Hz, 1H), 7.68 (t, *J* = 1.0 Hz, 1H), 6.91 – 6.87 (m, 1H), 6.84 (d, *J* = 8.7 Hz, 2H), 6.70 (d, *J* = 2.7 Hz, 2H), 6.32 (dd, *J* = 8.7, 2.7 Hz, 2H), 4.28 – 4.17 (m, 8H), 4.13 – 4.03 (m, 2H), 3.67 – 3.59 (m, 6H), 3.57 – 3.54 (m, 2H), 3.50 (t, *J* = 6.7 Hz, 2H), 3.40 (t, *J* = 6.7 Hz, 2H), 2.96 (s, 6H), 1.78 – 1.67 (m, 2H), 1.51 (p, *J* = 6.8 Hz, 2H), 1.44 – 1.35 (m, 2H), 1.35 – 1.26 (m, 2H), 0.65 (s, 3H), 0.57 (s, 3H); Analytical HPLC: $t_R$ = 13.0 min, 98% purity (10–95% MeCN/$H_2O$, linear gradient, with constant 0.1% v/v TFA additive, 20 min run, 1 mL/min flow, detection at 254 nm); HRMS (ESI) calculated for $C_{41}H_{53}ClN_3O_9S_2Si$ $[M+H]^+$ 858.2681, found 858.2690.

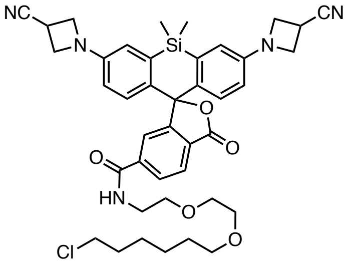


**(3-HTL; $JF_{629}$-HTL):** Synthesized following procedure B from **S3**. Purification by silica gel chromatography (20–100% EtOAc/hexanes, linear gradient) afforded **3-HTL** (73%) as a light blue solid. $^1H$ NMR ($CDCl_3$, 400 MHz) δ 7.98 (dd, *J* = 8.0, 0.7 Hz, 1H), 7.88 (dd, *J* = 8.0, 1.4 Hz, 1H), 7.70 (t, *J* = 1.0 Hz, 1H), 6.89 – 6.82 (m, 3H), 6.67 (d, *J* = 2.6 Hz, 2H), 6.30 (dd, *J* = 8.7, 2.7 Hz, 2H), 4.20 (dd, *J* = 8.5, 7.0 Hz, 4H), 4.09 (q, *J* = 6.7 Hz, 4H), 3.65 – 3.54 (m, 10H), 3.50 (t, *J* = 6.6 Hz, 2H), 3.41 (t, *J* = 6.7 Hz, 2H), 1.77 – 1.69 (m, 4H), 1.52 (p, *J* = 6.9 Hz, 2H), 1.44 – 1.36 (m, 2H), 1.35 – 1.28 (m, 2H), 0.66 (s, 3H), 0.58 (s, 3H); Analytical HPLC: $t_R$ = 14.4 min, 97% purity (10–95% MeCN/$H_2O$, linear gradient, with constant 0.1% v/v TFA additive, 20 min run, 1 mL/min flow, detection at 254 nm); HRMS (ESI) calculated for $C_{41}H_{47}ClN_5O_5Si$ $[M+H]^+$ 752.3035, found 752.3044.

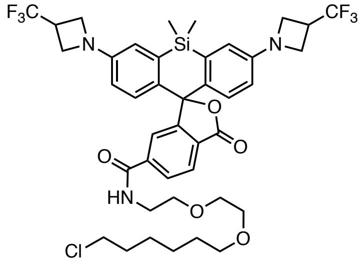


**(4-HTL; $JF_{626}$-HTL):** Synthesized following procedure B from **S4**. Purification by silica gel chromatography (30–100% EtOAc/hexanes, linear gradient) afforded **4-HTL** (83%) as a light blue solid. $^1H$ NMR ($CDCl_3$, 400 MHz) δ $^1H$ 7.99 (d, *J* = 7.8 Hz, 1H), 7.89 (dd, *J* = 8.0, 1.4 Hz, 1H), 7.73 – 7.68 (m, 1H), 6.84 – 6.78 (m, 3H), 6.67 (d, *J* = 2.7 Hz, 2H), 6.29 (dd, *J* = 8.7, 2.7 Hz, 2H), 4.07 (t, *J* = 8.1 Hz, 4H), 4.01 – 3.92 (m, 4H), 3.68 – 3.60 (m, 6H), 3.58 – 3.54 (m, 2H), 3.50 (t, *J* = 6.6 Hz, 2H), 3.45 – 3.34 (m, 4H), 1.77 – 1.68 (m, 2H), 1.56 – 1.46 (m, 2H), 1.44 – 1.35 (m, 2H), 1.34 – 1.26 (m, 2H), 0.65 (s, 3H), 0.58 (s, 3H); $^{19}F$ NMR ($CDCl_3$,

376 MHz) = -73.5 (d, $^3J_{HF}$ = 8.7 Hz); Analytical HPLC: $t_R$ = 16.4 min, 98% purity (10–95% MeCN/$H_2O$, linear gradient, with constant 0.1% v/v TFA additive, 20 min run, 1 mL/min flow, detection at 254 nm); HRMS (ESI) calculated for $C_{41}H_{47}ClF_6N_3O_5Si$ $[M+H]^+$ 838.2878, found 838.2891.

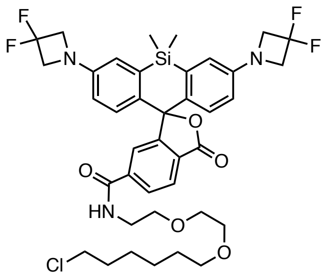


**(5-HTL; JF$_{614}$-HTL):** Synthesized following procedure B from **S5**. Purification by silica gel chromatography (0–3% MeOH/$CH_2Cl_2$, linear gradient) afforded **5-HTL** (75%) as an off-white solid. $^1H$ NMR ($CDCl_3$, 400 MHz) δ 8.00 (dd, *J* = 7.9, 0.7 Hz, 1H), 7.88 (dd, *J* = 8.0, 1.4 Hz, 1H), 7.70 (t, *J* = 1.0 Hz, 1H), 6.87 (d, *J* = 8.7 Hz, 2H), 6.77 – 6.70 (m, 3H), 6.36 (dd, *J* = 8.7, 2.7 Hz, 2H), 4.24 (t, *J* = 11.7 Hz, 8H), 3.67 – 3.59 (m, 6H), 3.58 – 3.53 (m, 2H), 3.50 (t, *J* = 6.6 Hz, 2H), 3.41 (t, *J* = 6.7 Hz, 2H), 1.78 – 1.69 (m, 2H), 1.57 – 1.49 (m, 2H), 1.44 – 1.36 (m, 2H), 1.35 – 1.28 (m, 2H), 0.67 (s, 3H), 0.60 (s, 3H); $^{19}F$ NMR ($CDCl_3$, 376 MHz) = - 99.9 (p, $^3J_{HF}$ = 11.6 Hz); Analytical HPLC: $t_R$ = 16.3 min, 95% purity (10–95% MeCN/$H_2O$, linear gradient, with constant 0.1% v/v TFA additive, 20 min run, 1 mL/min flow, detection at 254 nm); HRMS (ESI) calculated for $C_{39}H_{45}ClF_4N_3O_5Si$ $[M+H]^+$ 774.2753, found 774.2759.

### NMR SPECTRA AND HPLC TRACES

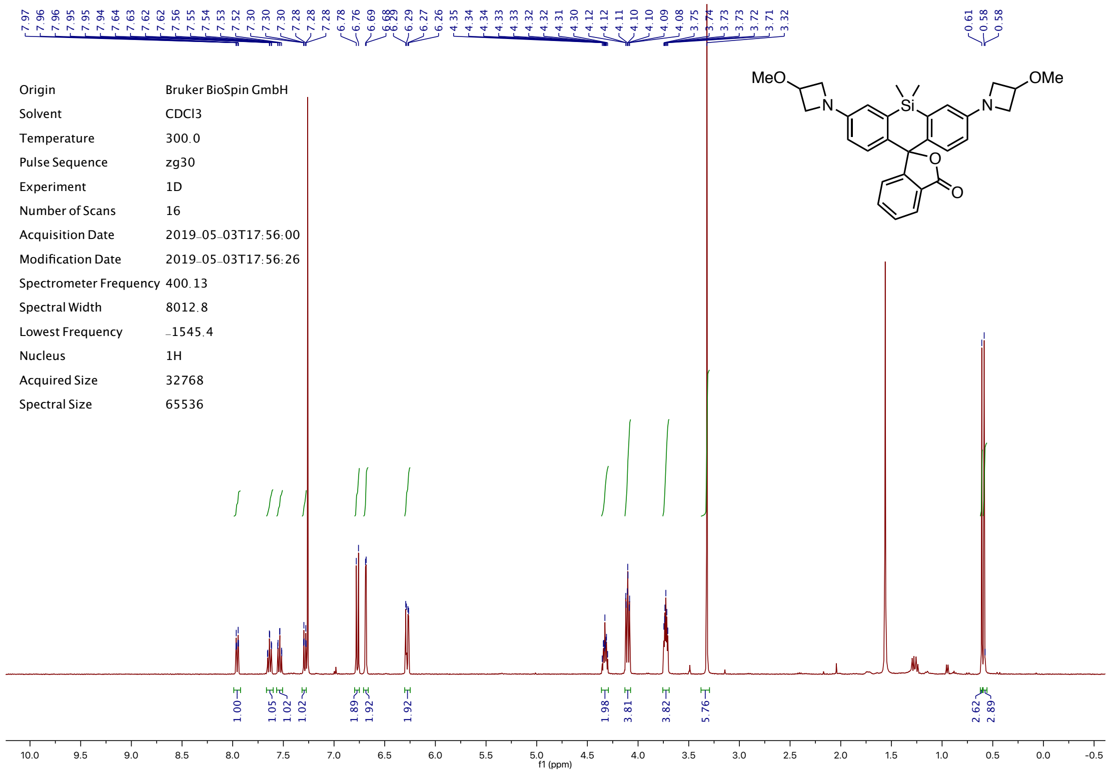


| | |
|---|---|
| Origin | Bruker BioSpin GmbH |
| Solvent | CDCl3 |
| Temperature | 300.0 |
| Pulse Sequence | zg30 |
| Experiment | 1D |
| Number of Scans | 16 |
| Acquisition Date | 2019-05-03T17:56:00 |
| Modification Date | 2019-05-03T17:56:26 |
| Spectrometer Frequency | 400.13 |
| Spectral Width | 8012.8 |
| Lowest Frequency | -1545.4 |
| Nucleus | 1H |
| Acquired Size | 32768 |
| Spectral Size | 65536 |

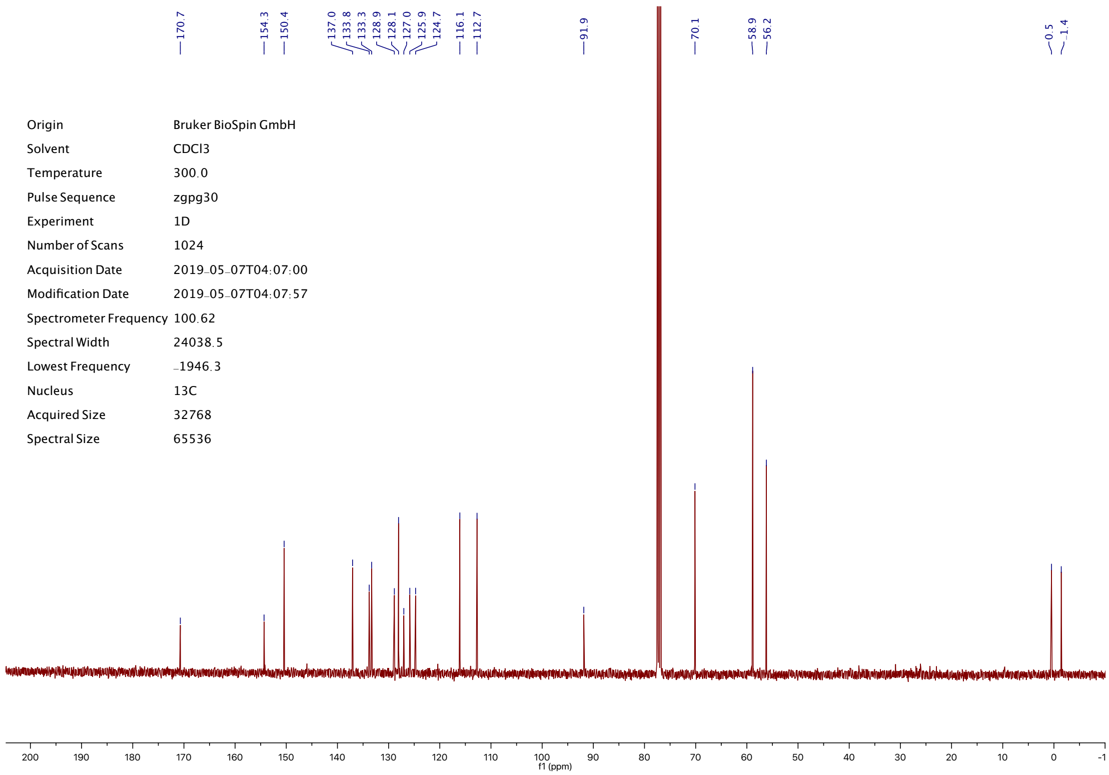


| | |
|---|---|
| Origin | Bruker BioSpin GmbH |
| Solvent | CDCl3 |
| Temperature | 300.0 |
| Pulse Sequence | zgpg30 |
| Experiment | 1D |
| Number of Scans | 1024 |
| Acquisition Date | 2019-05-07T04:07:00 |
| Modification Date | 2019-05-07T04:07:57 |
| Spectrometer Frequency | 100.62 |
| Spectral Width | 24038.5 |
| Lowest Frequency | -1946.3 |
| Nucleus | 13C |
| Acquired Size | 32768 |
| Spectral Size | 65536 |

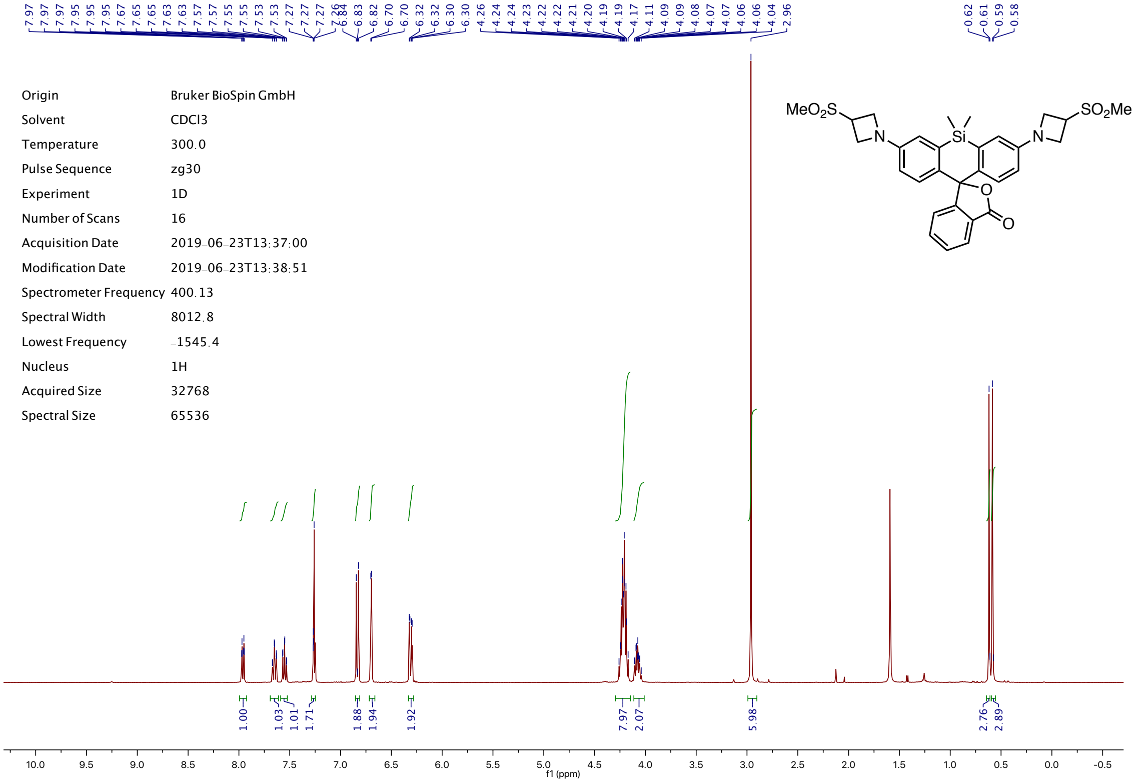


| | |
|---|---|
| Origin | Bruker BioSpin GmbH |
| Solvent | CDCl3 |
| Temperature | 300.0 |
| Pulse Sequence | zg30 |
| Experiment | 1D |
| Number of Scans | 16 |
| Acquisition Date | 2019-06-23T13:37:00 |
| Modification Date | 2019-06-23T13:38:51 |
| Spectrometer Frequency | 400.13 |
| Spectral Width | 8012.8 |
| Lowest Frequency | -1545.4 |
| Nucleus | 1H |
| Acquired Size | 32768 |
| Spectral Size | 65536 |



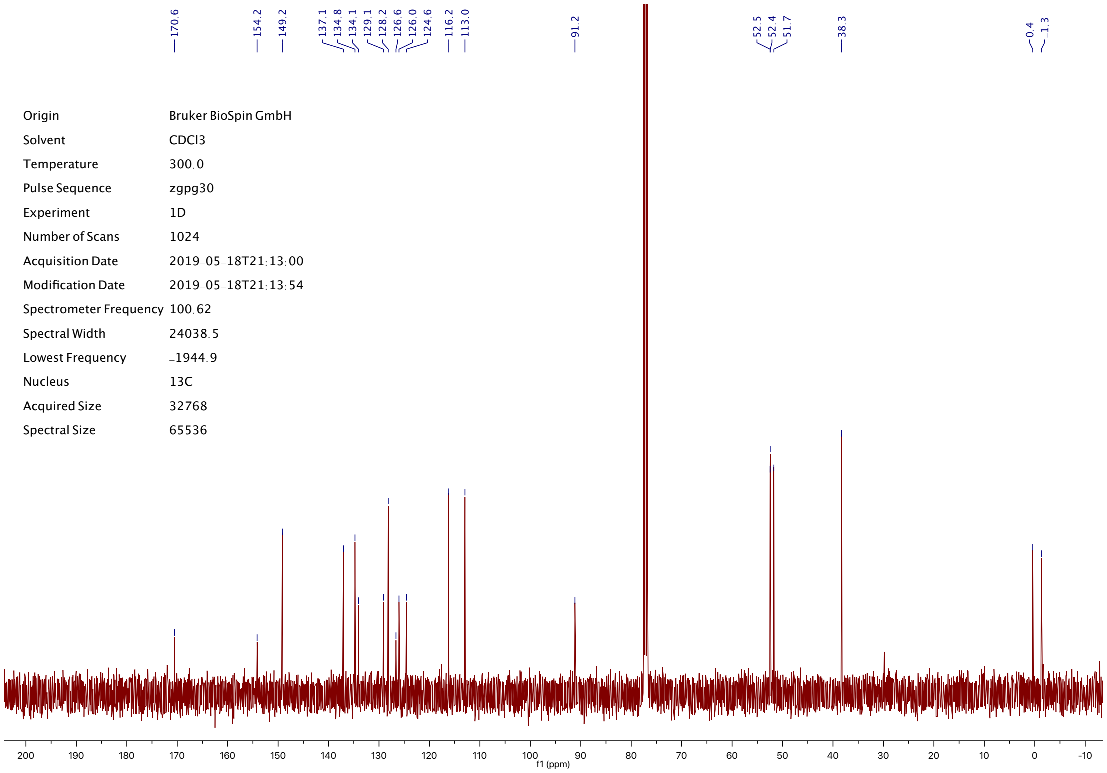


| | |
|---|---|
| Origin | Bruker BioSpin GmbH |
| Solvent | CDCl3 |
| Temperature | 300.0 |
| Pulse Sequence | zgpg30 |
| Experiment | 1D |
| Number of Scans | 1024 |
| Acquisition Date | 2019-05-18T21:13:00 |
| Modification Date | 2019-05-18T21:13:54 |
| Spectrometer Frequency | 100.62 |
| Spectral Width | 24038.5 |
| Lowest Frequency | -1944.9 |
| Nucleus | 13C |
| Acquired Size | 32768 |
| Spectral Size | 65536 |

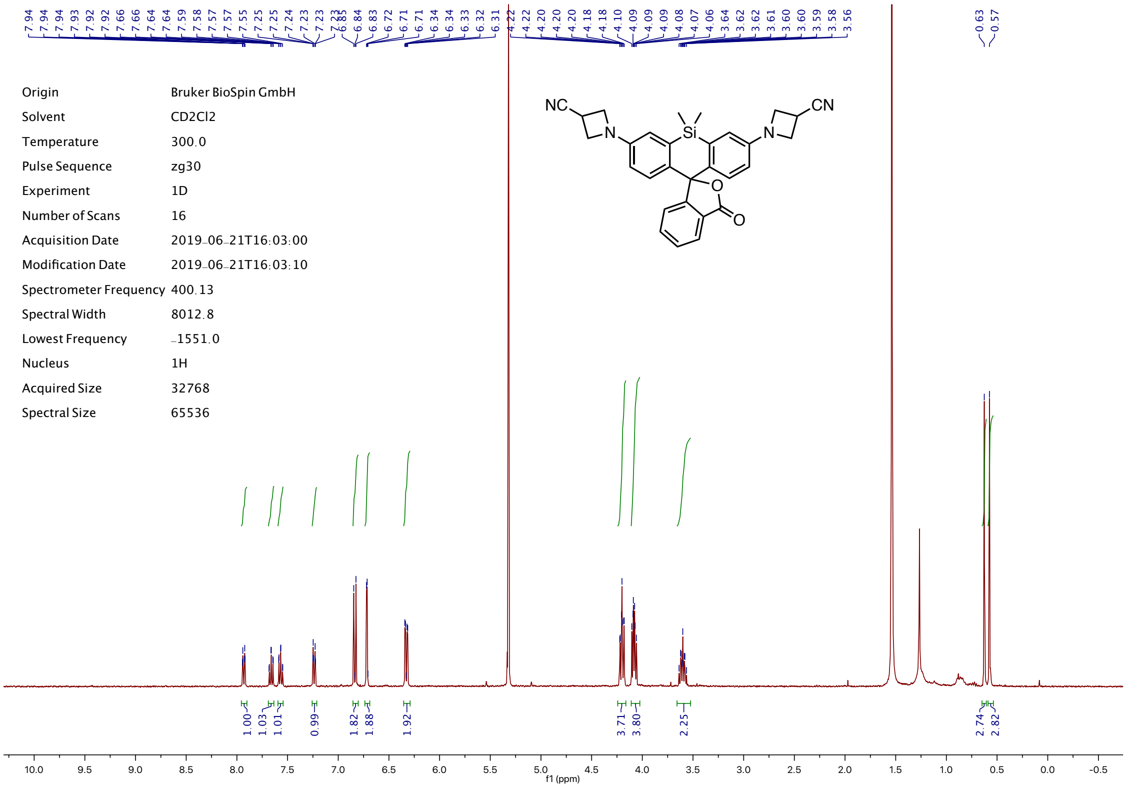


| | |
|---|---|
| Origin | Bruker BioSpin GmbH |
| Solvent | CD2Cl2 |
| Temperature | 300.0 |
| Pulse Sequence | zg30 |
| Experiment | 1D |
| Number of Scans | 16 |
| Acquisition Date | 2019-06-21T16:03:00 |
| Modification Date | 2019-06-21T16:03:10 |
| Spectrometer Frequency | 400.13 |
| Spectral Width | 8012.8 |
| Lowest Frequency | -1551.0 |
| Nucleus | 1H |
| Acquired Size | 32768 |
| Spectral Size | 65536 |

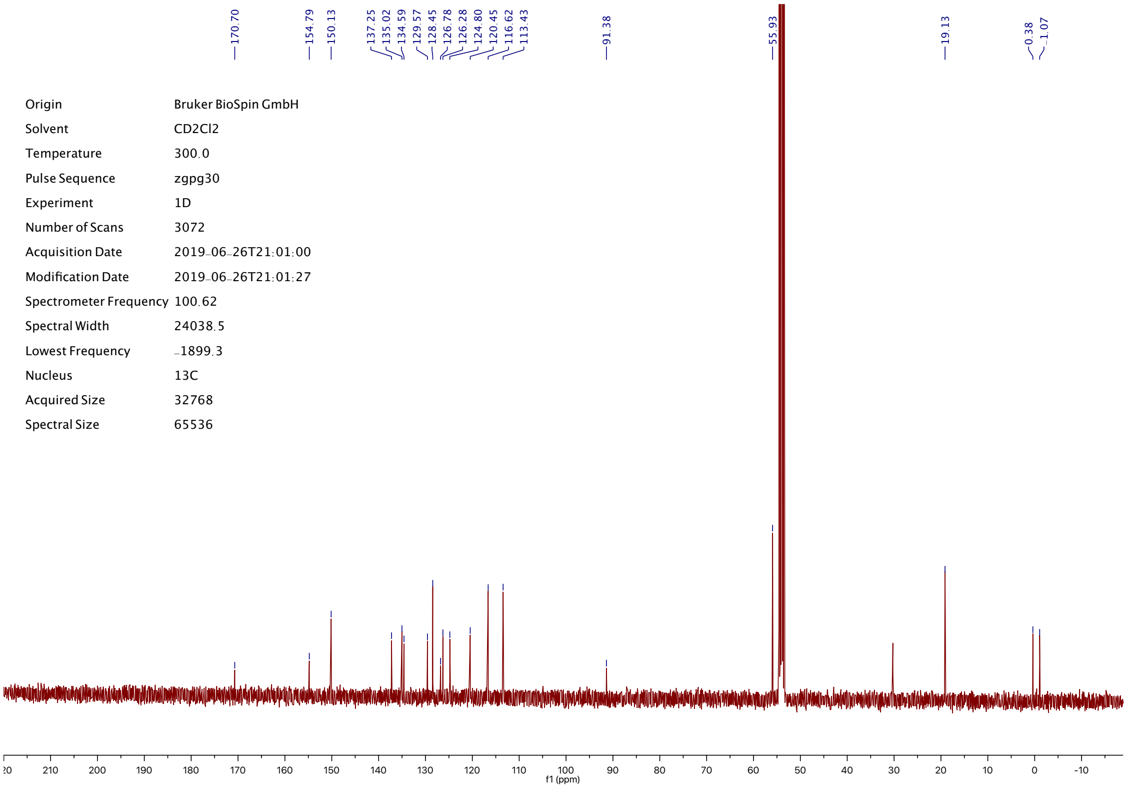


| | |
|---|---|
| Origin | Bruker BioSpin GmbH |
| Solvent | CD2Cl2 |
| Temperature | 300.0 |
| Pulse Sequence | zgpg30 |
| Experiment | 1D |
| Number of Scans | 3072 |
| Acquisition Date | 2019-06-26T21:01:00 |
| Modification Date | 2019-06-26T21:01:27 |
| Spectrometer Frequency | 100.62 |
| Spectral Width | 24038.5 |
| Lowest Frequency | -1899.3 |
| Nucleus | 13C |
| Acquired Size | 32768 |
| Spectral Size | 65536 |

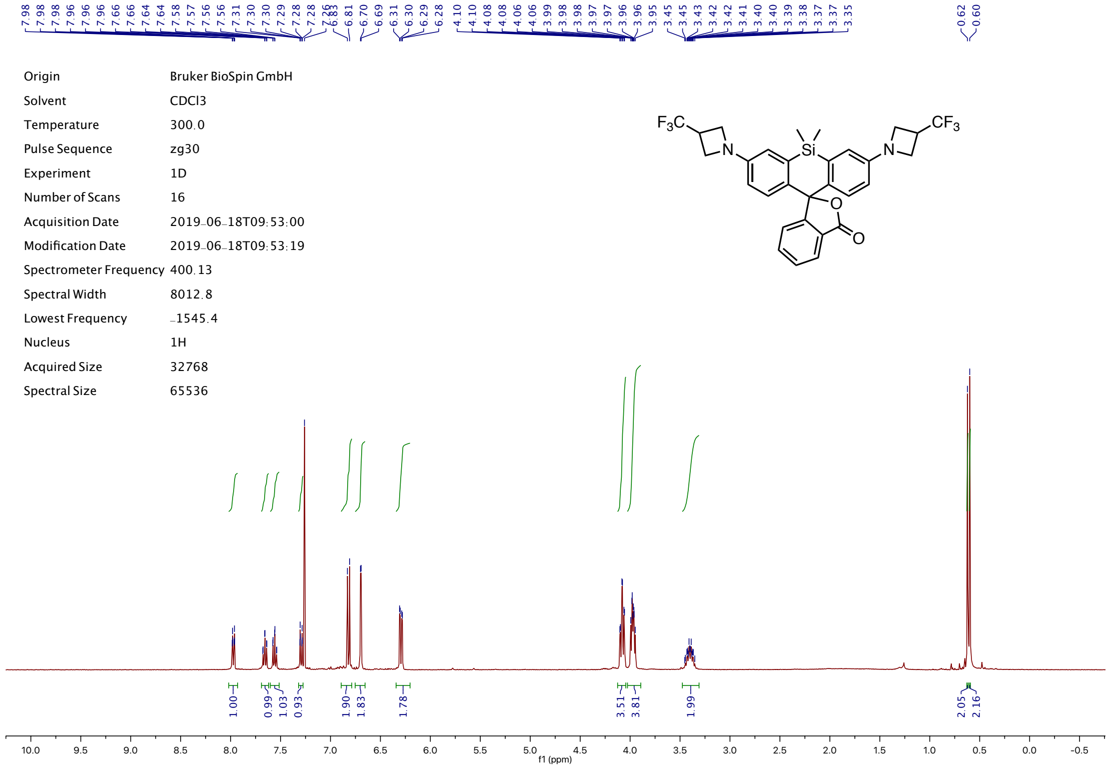
Origin Bruker BioSpin GmbH
Solvent CDCl3
Temperature 300.0
Pulse Sequence zg30
Experiment 1D
Number of Scans 16
Acquisition Date 2019-06-18T09:53:00
Modification Date 2019-06-18T09:53:19
Spectrometer Frequency 400.13
Spectral Width 8012.8
Lowest Frequency -1545.4
Nucleus 1H
Acquired Size 32768
Spectral Size 65536
f1 (ppm)

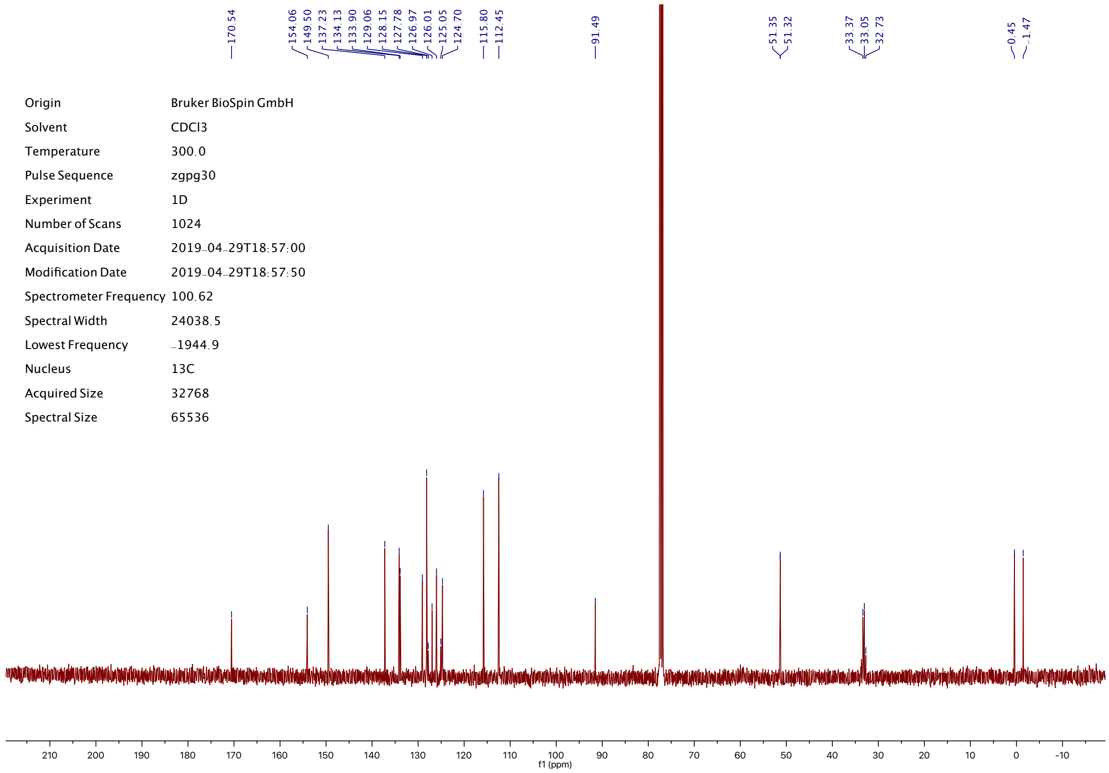
Origin Bruker BioSpin GmbH
Solvent CDCl3
Temperature 300.0
Pulse Sequence zgpg30
Experiment 1D
Number of Scans 1024
Acquisition Date 2019-04-29T18:57:00
Modification Date 2019-04-29T18:57:50
Spectrometer Frequency 100.62
Spectral Width 24038.5
Lowest Frequency -1944.9
Nucleus 13C
Acquired Size 32768
Spectral Size 65536
170.54
154.06
149.50
137.23
134.13
133.90
129.06
128.15
127.78
126.97
126.01
125.05
124.70
115.80
112.45
91.49
51.35
51.32
33.37
33.05
32.73
0.45
-1.47
f1 (ppm)

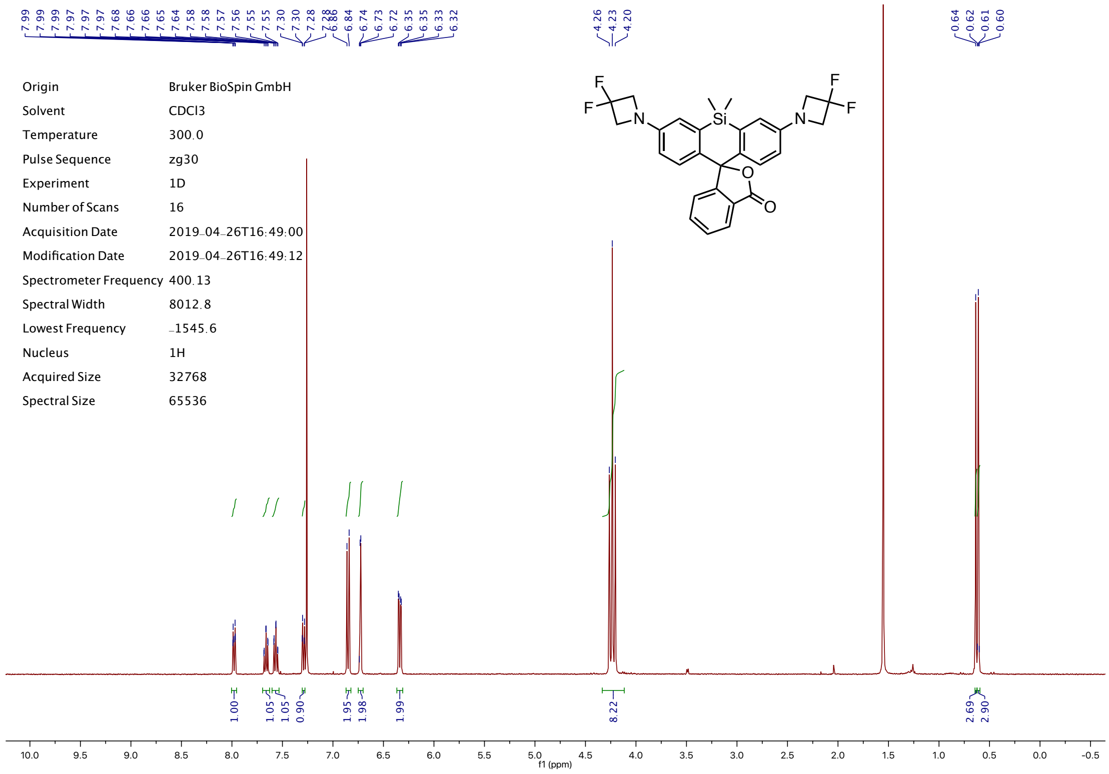
Origin Bruker BioSpin GmbH
Solvent CDCl3
Temperature 300.0
Pulse Sequence zg30
Experiment 1D
Number of Scans 16
Acquisition Date 2019-04-26T16:49:00
Modification Date 2019-04-26T16:49:12
Spectrometer Frequency 400.13
Spectral Width 8012.8
Lowest Frequency -1545.6
Nucleus 1H
Acquired Size 32768
Spectral Size 65536
4.26
4.23
4.20
0.64
0.62
0.61
0.60
1.00
1.05
1.05
0.90
1.95
1.98
1.99
8.22
2.69
2.90
f1 (ppm)

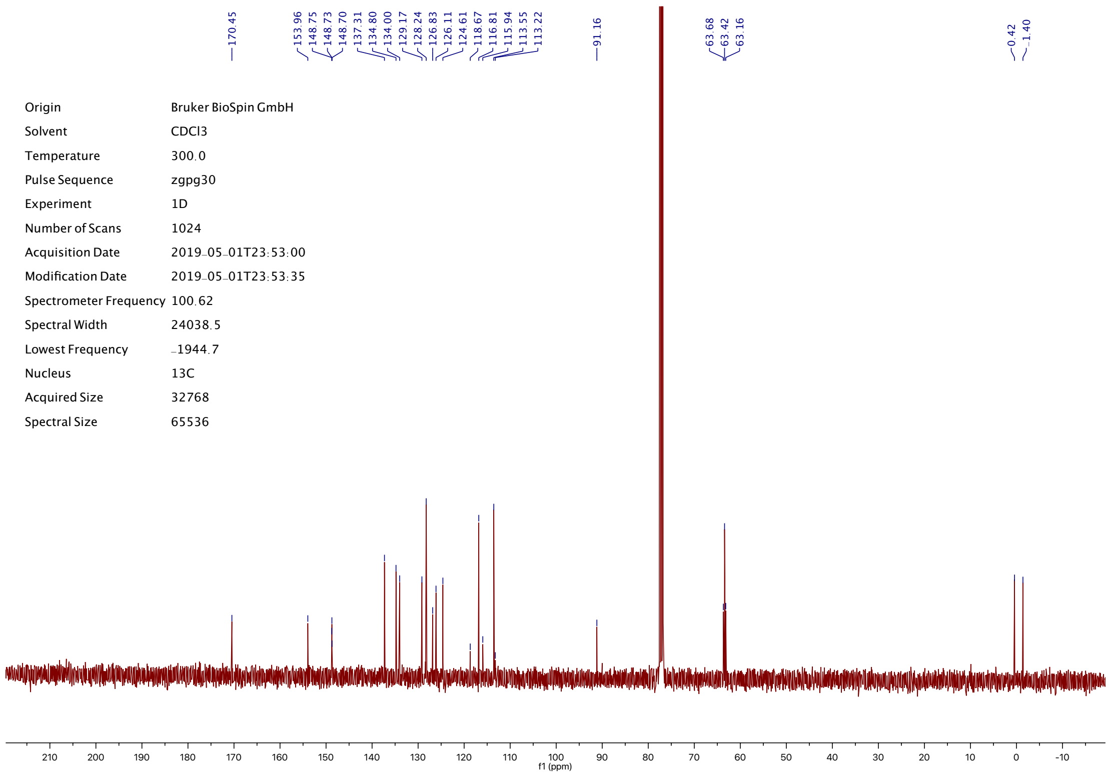
170.45
153.96
148.75
148.73
148.70
137.31
134.80
134.00
129.17
128.24
126.83
126.11
124.61
118.67
116.81
115.94
113.55
113.22
91.16
63.68
63.42
63.16
0.42
-1.40
Origin Bruker BioSpin GmbH
Solvent CDCl3
Temperature 300.0
Pulse Sequence zgpg30
Experiment 1D
Number of Scans 1024
Acquisition Date 2019-05-01T23:53:00
Modification Date 2019-05-01T23:53:35
Spectrometer Frequency 100.62
Spectral Width 24038.5
Lowest Frequency -1944.7
Nucleus 13C
Acquired Size 32768
Spectral Size 65536
f1 (ppm)

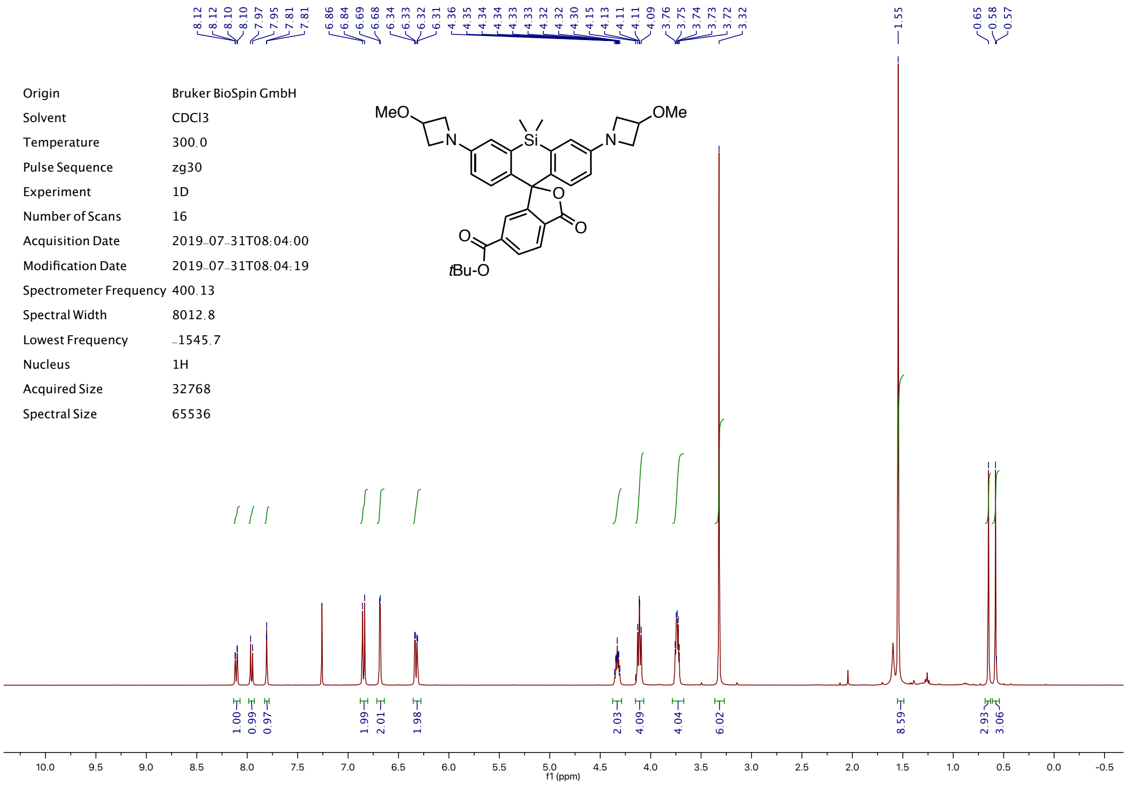

8.12
8.12
8.10
8.10
7.97
7.95
7.81
7.81
6.86
6.84
6.69
6.68
6.34
6.33
6.32
6.31
4.36
4.35
4.34
4.34
4.33
4.33
4.32
4.32
4.30
4.15
4.13
4.11
4.11
4.09
3.76
3.75
3.74
3.73
3.73
3.72
3.32
1.55
0.65
0.58
0.57
Origin Bruker BioSpin GmbH
Solvent CDCl3
Temperature 300.0
Pulse Sequence zg30
Experiment 1D
Number of Scans 16
Acquisition Date 2019-07-31T08:04:00
Modification Date 2019-07-31T08:04:19
Spectrometer Frequency 400.13
Spectral Width 8012.8
Lowest Frequency -1545.7
Nucleus 1H
Acquired Size 32768
Spectral Size 65536
MeO
OMe
N
Si
O
tBu-O
1.00
0.99
0.97
1.99
2.01
1.98
2.03
4.09
4.04
6.02
8.59
2.93
3.06
10.0 9.5 9.0 8.5 8.0 7.5 7.0 6.5 6.0 5.5 5.0 4.5 4.0 3.5 3.0 2.5 2.0 1.5 1.0 0.5 0.0 -0.5
f1 (ppm)


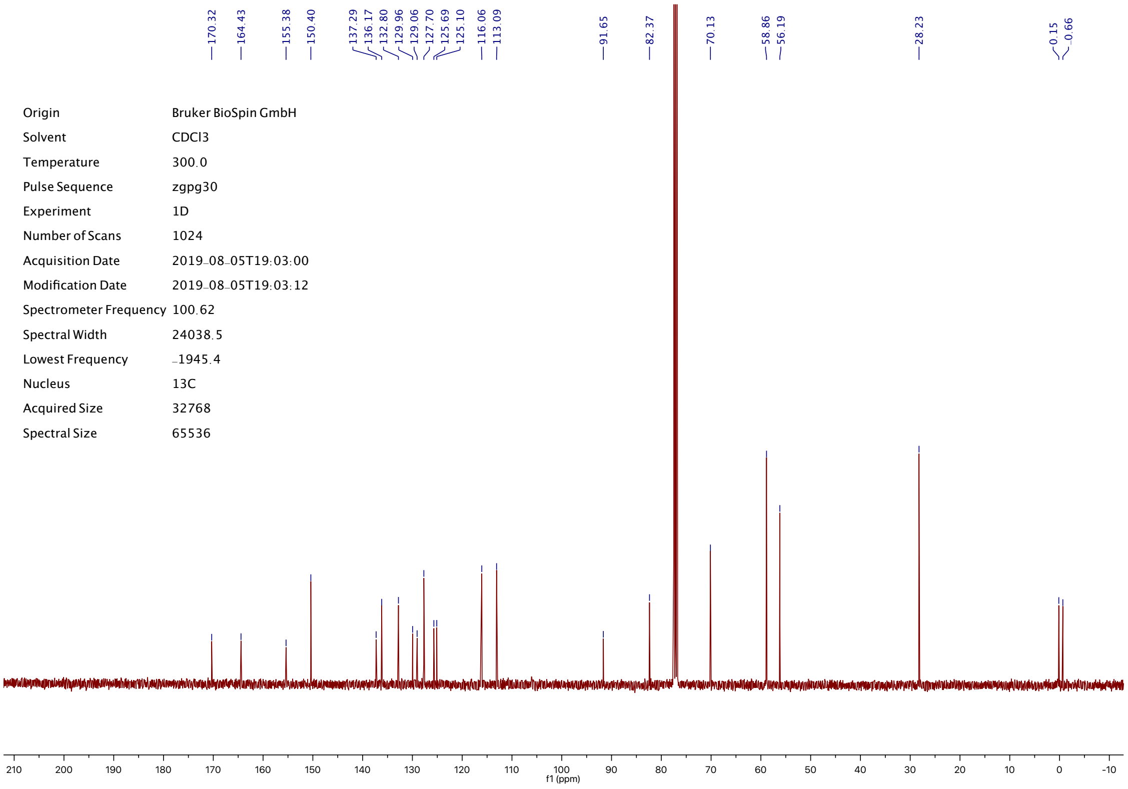

170.32
164.43
155.38
150.40
137.29
136.17
132.80
129.96
129.06
127.70
125.69
125.10
116.06
113.09
91.65
82.37
70.13
58.86
56.19
28.23
0.15
-0.66
Origin Bruker BioSpin GmbH
Solvent CDCl3
Temperature 300.0
Pulse Sequence zgpg30
Experiment 1D
Number of Scans 1024
Acquisition Date 2019-08-05T19:03:00
Modification Date 2019-08-05T19:03:12
Spectrometer Frequency 100.62
Spectral Width 24038.5
Lowest Frequency -1945.4
Nucleus 13C
Acquired Size 32768
Spectral Size 65536
210 200 190 180 170 160 150 140 130 120 110 100 90 80 70 60 50 40 30 20 10 0 -10
f1 (ppm)

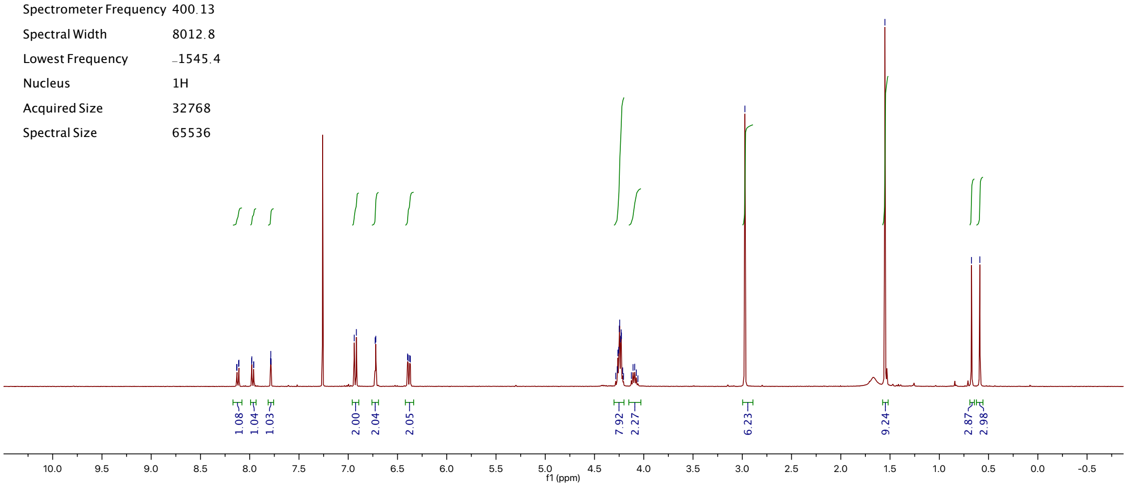


| | |
|---|---|
| Origin | Bruker BioSpin GmbH |
| Solvent | CDCl3 |
| Temperature | 300.0 |
| Pulse Sequence | zg30 |
| Experiment | 1D |
| Number of Scans | 16 |
| Acquisition Date | 2019-08-23T10:58:00 |
| Modification Date | 2019-08-23T10:58:39 |
| Spectrometer Frequency | 400.13 |
| Spectral Width | 8012.8 |
| Lowest Frequency | -1545.4 |
| Nucleus | 1H |
| Acquired Size | 32768 |
| Spectral Size | 65536 |

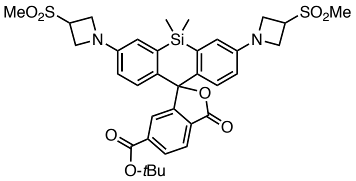


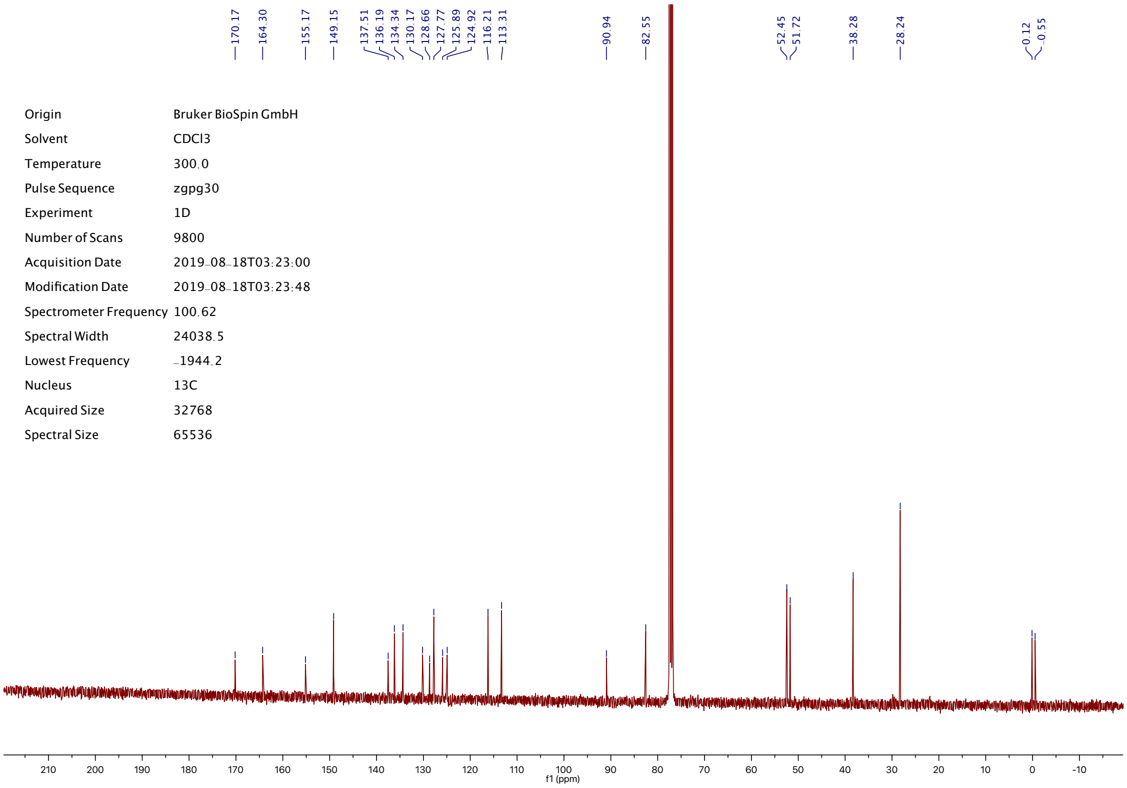


| | |
|---|---|
| Origin | Bruker BioSpin GmbH |
| Solvent | CDCl3 |
| Temperature | 300.0 |
| Pulse Sequence | zgpg30 |
| Experiment | 1D |
| Number of Scans | 9800 |
| Acquisition Date | 2019-08-18T03:23:00 |
| Modification Date | 2019-08-18T03:23:48 |
| Spectrometer Frequency | 100.62 |
| Spectral Width | 24038.5 |
| Lowest Frequency | -1944.2 |
| Nucleus | 13C |
| Acquired Size | 32768 |
| Spectral Size | 65536 |

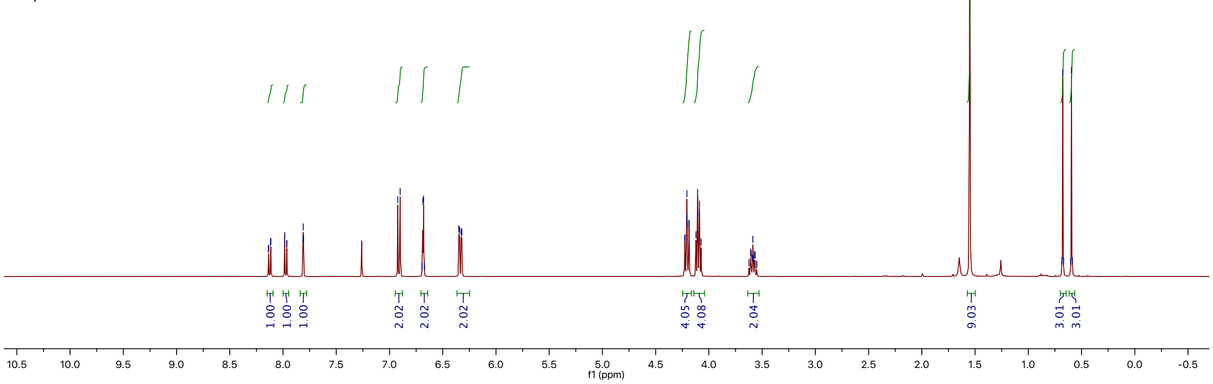


| | |
|---|---|
| Origin | Bruker BioSpin GmbH |
| Solvent | CDCl3 |
| Temperature | 300.0 |
| Pulse Sequence | zg30 |
| Experiment | 1D |
| Number of Scans | 16 |
| Acquisition Date | 2019-08-07T10:40:00 |
| Modification Date | 2019-08-07T10:40:26 |
| Spectrometer Frequency | 400.13 |
| Spectral Width | 8012.8 |
| Lowest Frequency | -1545.4 |
| Nucleus | 1H |
| Acquired Size | 32768 |
| Spectral Size | 65536 |

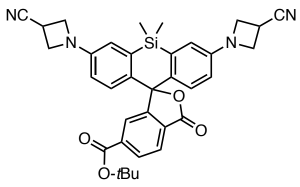


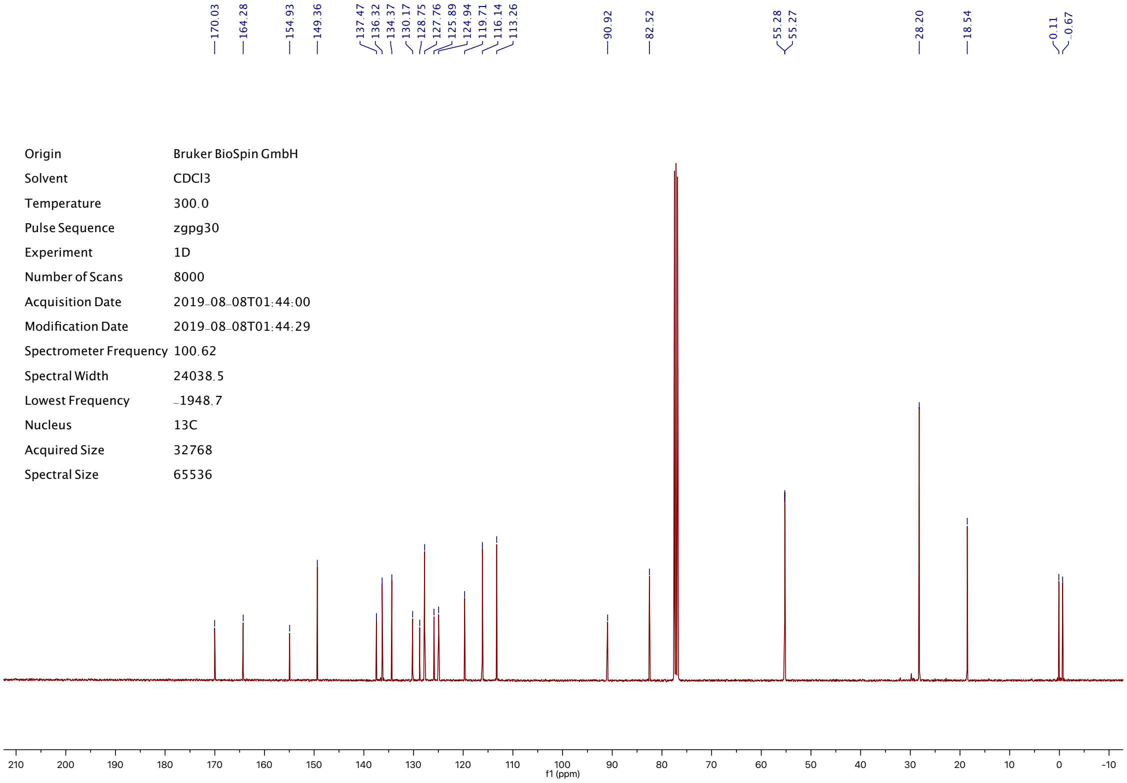


| | |
|---|---|
| Origin | Bruker BioSpin GmbH |
| Solvent | CDCl3 |
| Temperature | 300.0 |
| Pulse Sequence | zgpg30 |
| Experiment | 1D |
| Number of Scans | 8000 |
| Acquisition Date | 2019-08-08T01:44:00 |
| Modification Date | 2019-08-08T01:44:29 |
| Spectrometer Frequency | 100.62 |
| Spectral Width | 24038.5 |
| Lowest Frequency | -1948.7 |
| Nucleus | 13C |
| Acquired Size | 32768 |
| Spectral Size | 65536 |

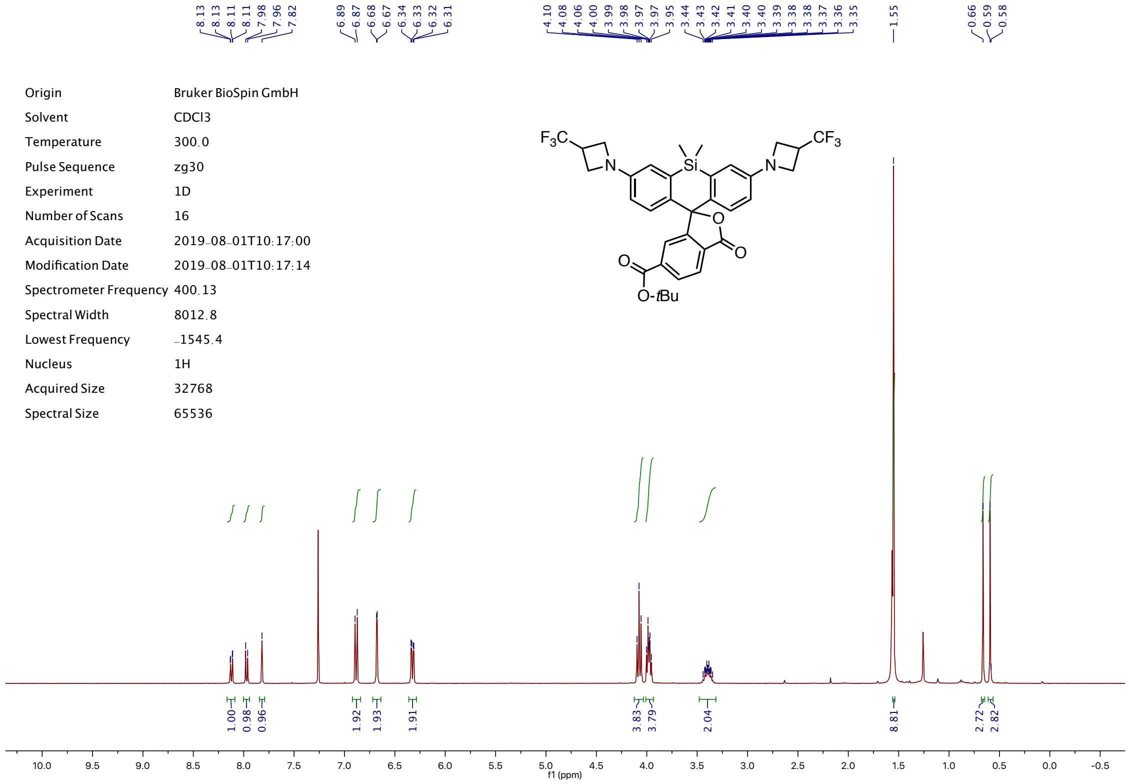
8.13
8.13
8.11
8.11
7.98
7.96
7.82
6.89
6.87
6.68
6.67
6.34
6.33
6.32
6.31
4.10
4.08
4.06
4.00
3.99
3.98
3.97
3.97
3.95
3.44
3.43
3.42
3.41
3.40
3.40
3.39
3.38
3.38
3.37
3.36
3.35
1.55
0.66
0.59
0.58
Origin Bruker BioSpin GmbH
Solvent CDCl3
Temperature 300.0
Pulse Sequence zg30
Experiment 1D
Number of Scans 16
Acquisition Date 2019-08-01T10:17:00
Modification Date 2019-08-01T10:17:14
Spectrometer Frequency 400.13
Spectral Width 8012.8
Lowest Frequency -1545.4
Nucleus 1H
Acquired Size 32768
Spectral Size 65536
F3C
N
Si
N
CF3
O
O
O
O-tBu
1.00
0.98
0.96
1.92
1.93
1.91
3.83
3.79
2.04
8.81
2.72
2.82
10.0 9.5 9.0 8.5 8.0 7.5 7.0 6.5 6.0 5.5 5.0 4.5 4.0 3.5 3.0 2.5 2.0 1.5 1.0 0.5 0.0 -0.5
f1 (ppm)

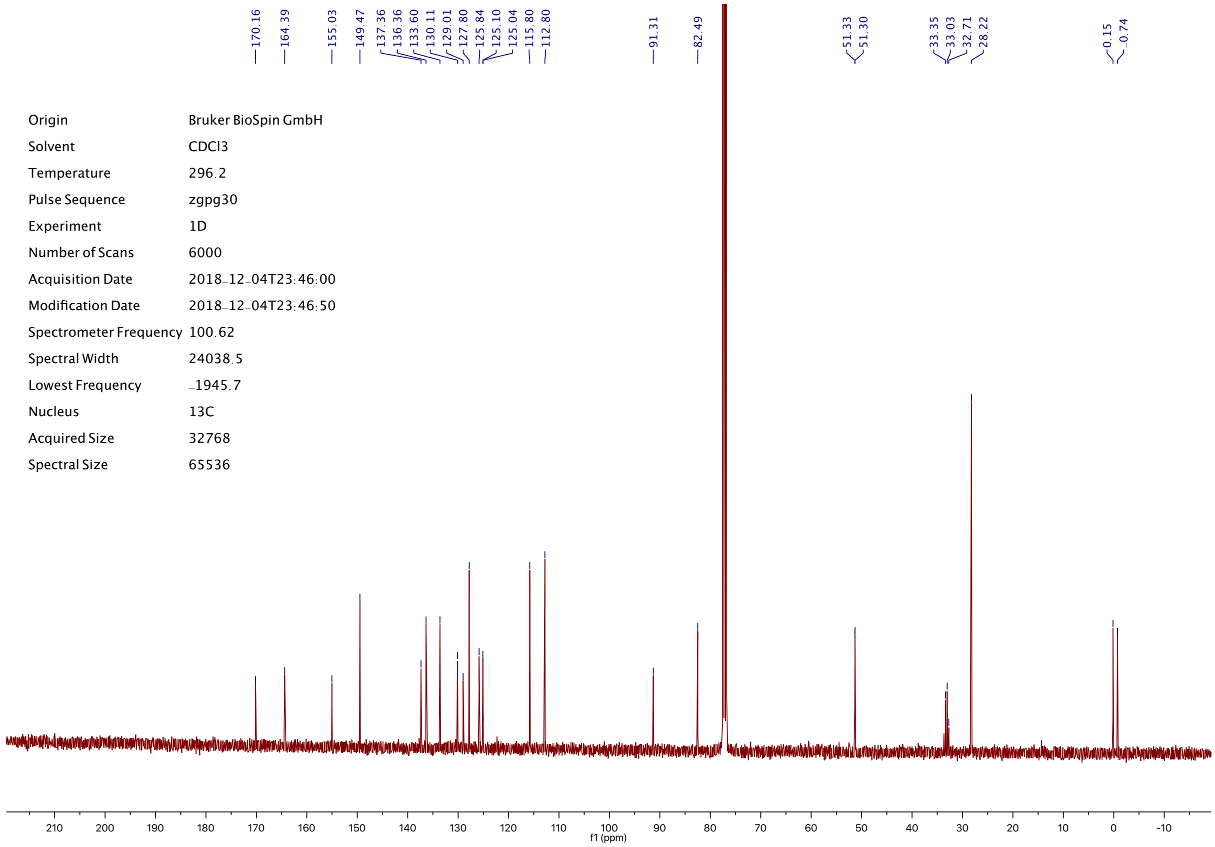
170.16
164.39
155.03
149.47
137.36
136.36
133.60
130.11
129.01
127.80
125.84
125.10
125.04
115.80
112.80
91.31
82.49
51.33
51.30
33.35
33.03
32.71
28.22
-0.15
-0.74
Origin Bruker BioSpin GmbH
Solvent CDCl3
Temperature 296.2
Pulse Sequence zgpg30
Experiment 1D
Number of Scans 6000
Acquisition Date 2018-12-04T23:46:00
Modification Date 2018-12-04T23:46:50
Spectrometer Frequency 100.62
Spectral Width 24038.5
Lowest Frequency -1945.7
Nucleus 13C
Acquired Size 32768
Spectral Size 65536
210 200 190 180 170 160 150 140 130 120 110 100 90 80 70 60 50 40 30 20 10 0 -10
f1 (ppm)

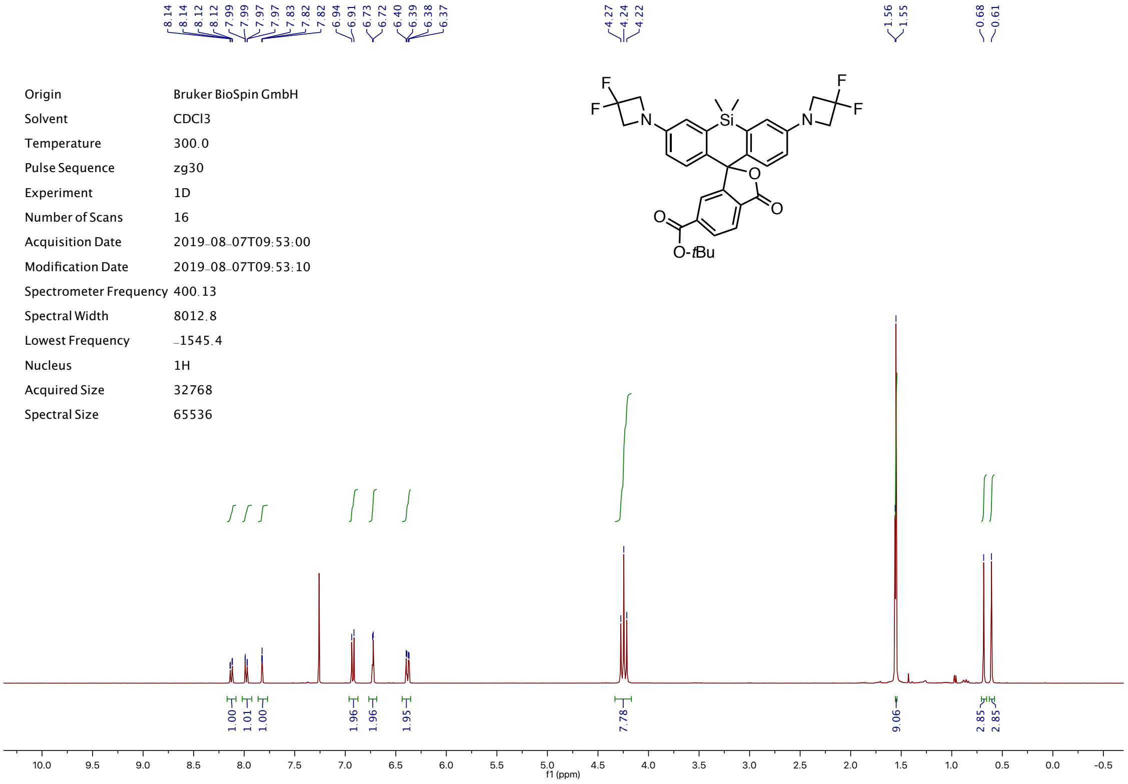
8.14
8.14
8.12
8.12
7.99
7.99
7.97
7.97
7.83
7.82
7.82
6.94
6.91
6.73
6.72
6.40
6.39
6.38
6.37
4.27
4.24
4.22
1.56
1.55
0.68
0.61
F
F
N
Si
N
F
F
O
O
O
O-tBu
Origin Bruker BioSpin GmbH
Solvent CDCl3
Temperature 300.0
Pulse Sequence zg30
Experiment 1D
Number of Scans 16
Acquisition Date 2019-08-07T09:53:00
Modification Date 2019-08-07T09:53:10
Spectrometer Frequency 400.13
Spectral Width 8012.8
Lowest Frequency -1545.4
Nucleus 1H
Acquired Size 32768
Spectral Size 65536
1.00
1.01
1.00
1.96
1.96
1.95
7.78
9.06
2.85
2.85
10.0 9.5 9.0 8.5 8.0 7.5 7.0 6.5 6.0 5.5 5.0 4.5 4.0 3.5 3.0 2.5 2.0 1.5 1.0 0.5 0.0 -0.5
f1 (ppm)

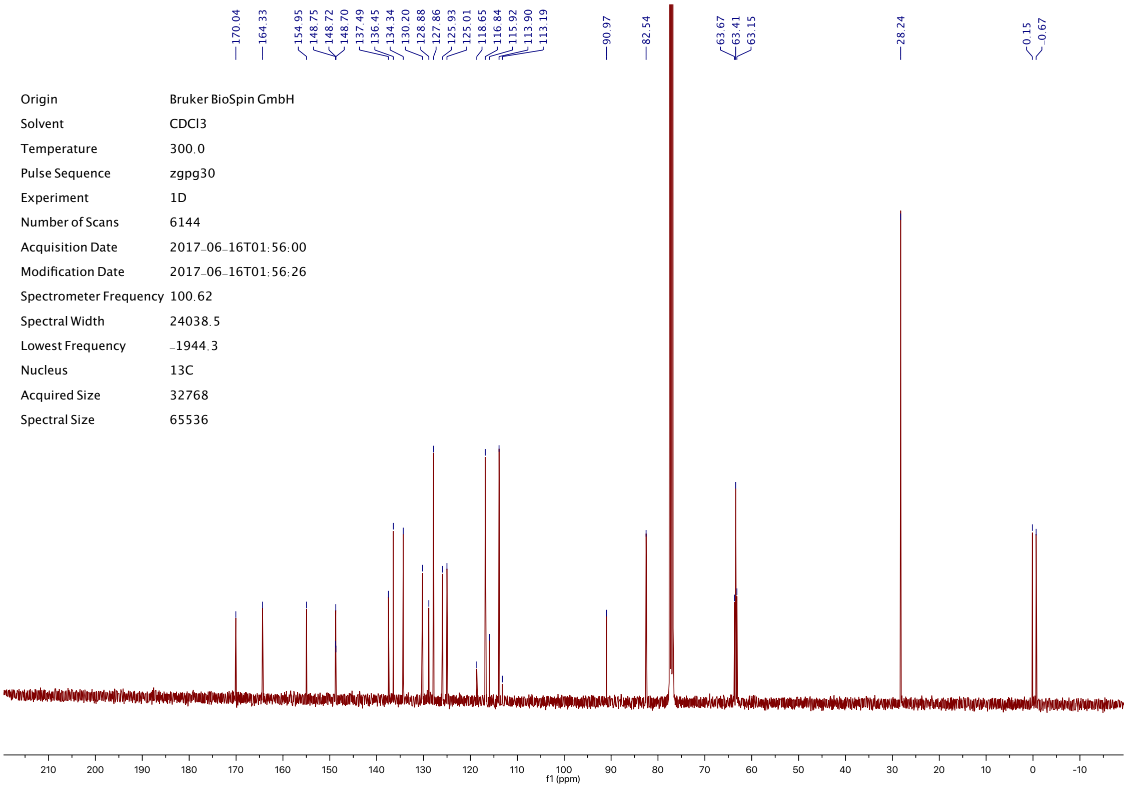
170.04
164.33
154.95
148.75
148.72
148.70
137.49
136.45
134.34
130.20
128.88
127.86
125.93
125.01
118.65
116.84
115.92
113.90
113.19
90.97
82.54
63.67
63.41
63.15
28.24
0.15
-0.67
Origin Bruker BioSpin GmbH
Solvent CDCl3
Temperature 300.0
Pulse Sequence zgpg30
Experiment 1D
Number of Scans 6144
Acquisition Date 2017-06-16T01:56:00
Modification Date 2017-06-16T01:56:26
Spectrometer Frequency 100.62
Spectral Width 24038.5
Lowest Frequency -1944.3
Nucleus 13C
Acquired Size 32768
Spectral Size 65536
210 200 190 180 170 160 150 140 130 120 110 100 90 80 70 60 50 40 30 20 10 0 -10
f1 (ppm)

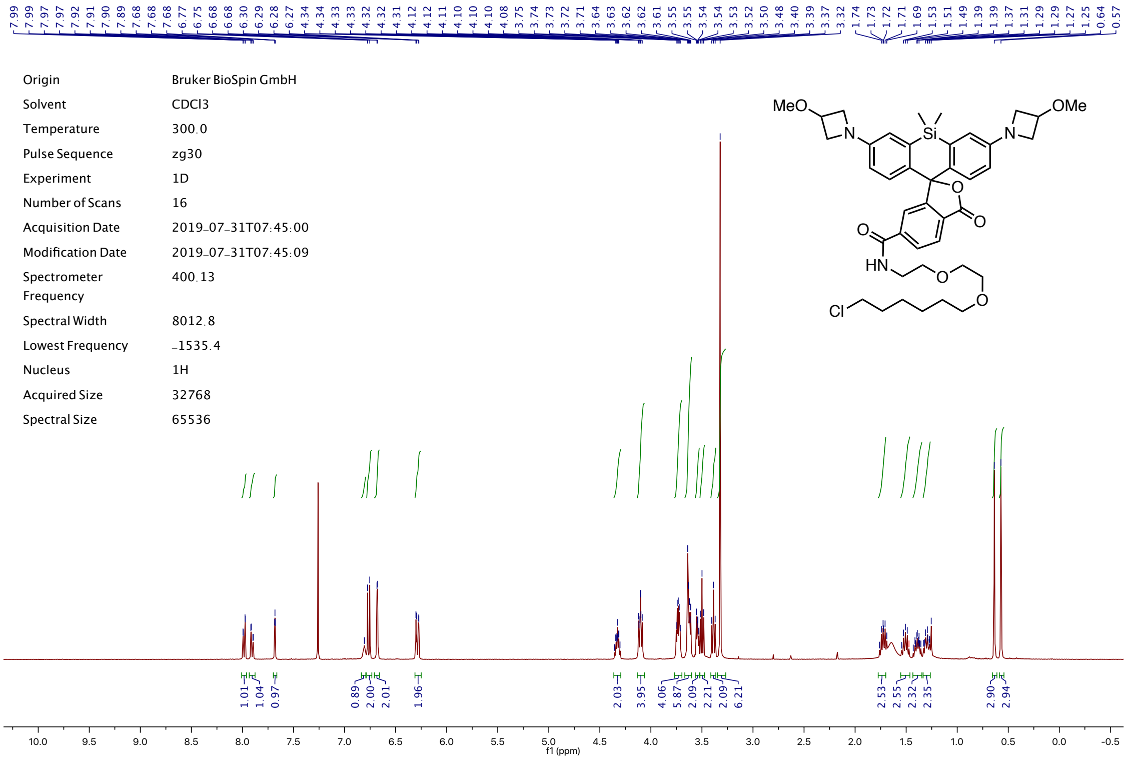
7.99 7.99 7.97 7.97 7.92 7.91 7.90 7.89 7.68 7.68 7.68 6.77 6.75 6.68 6.68 6.30 6.29 6.28 6.27 4.34 4.34 4.33 4.33 4.32 4.32 4.31 4.12 4.12 4.11 4.10 4.10 4.10 4.08 3.75 3.74 3.73 3.72 3.71 3.64 3.63 3.62 3.61 3.55 3.55 3.54 3.54 3.54 3.53 3.52 3.50 3.48 3.40 3.39 3.37 3.32 1.74 1.73 1.72 1.71 1.69 1.53 1.51 1.49 1.39 1.39 1.37 1.31 1.29 1.29 1.27 1.25 0.64 0.57
Origin Bruker BioSpin GmbH
Solvent CDCl3
Temperature 300.0
Pulse Sequence zg30
Experiment 1D
Number of Scans 16
Acquisition Date 2019-07-31T07:45:00
Modification Date 2019-07-31T07:45:09
Spectrometer Frequency 400.13
Spectral Width 8012.8
Lowest Frequency -1535.4
Nucleus 1H
Acquired Size 32768
Spectral Size 65536
MeO
OMe
Si
N
N
O
O
O
HN
O
O
Cl
1.01 1.04 0.97 0.89 2.00 2.01 1.96 2.03 3.95 4.06 5.87 2.09 2.21 2.09 6.21 2.53 2.55 2.32 2.35 2.90 2.94
10.0 9.5 9.0 8.5 8.0 7.5 7.0 6.5 6.0 5.5 5.0 4.5 4.0 3.5 3.0 2.5 2.0 1.5 1.0 0.5 0.0 -0.5
f1 (ppm)

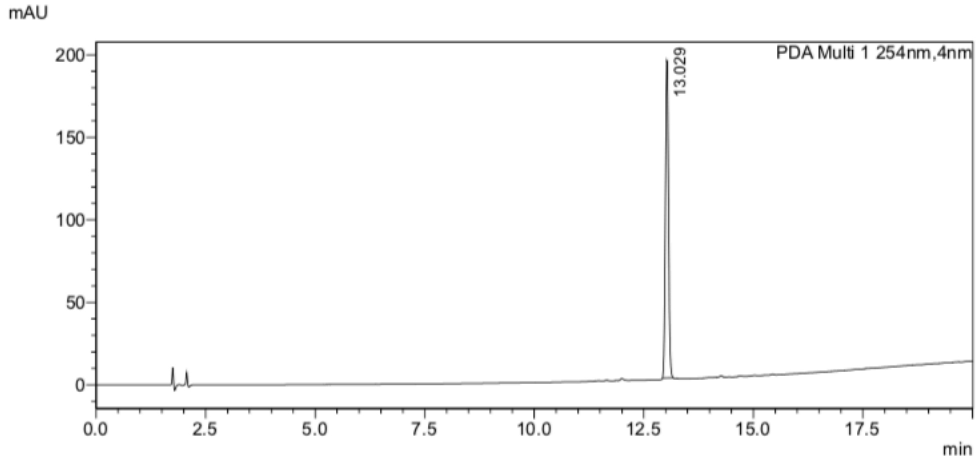
mAU
PDA Multi 1 254nm,4nm
13.029
200
150
100
50
0
0.0 2.5 5.0 7.5 10.0 12.5 15.0 17.5
min

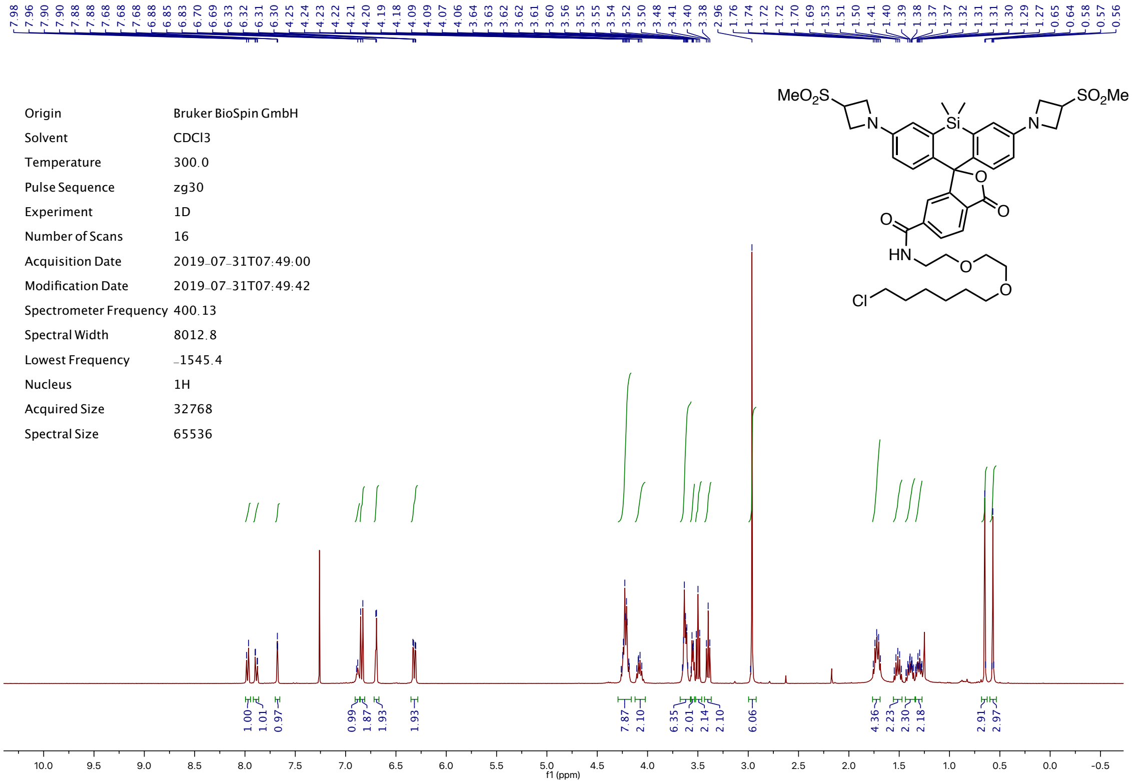
Origin Bruker BioSpin GmbH
Solvent CDCl3
Temperature 300.0
Pulse Sequence zg30
Experiment 1D
Number of Scans 16
Acquisition Date 2019-07-31T07:49:00
Modification Date 2019-07-31T07:49:42
Spectrometer Frequency 400.13
Spectral Width 8012.8
Lowest Frequency -1545.4
Nucleus 1H
Acquired Size 32768
Spectral Size 65536
f1 (ppm)
MeO2S
Si
N
O
HN
Cl
SO2Me

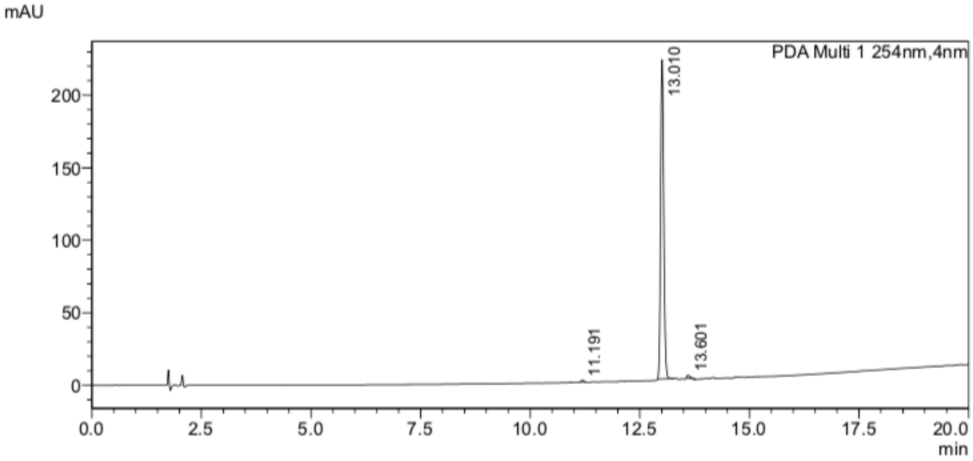
mAU
PDA Multi 1 254nm,4nm
13.010
11.191
13.601
min

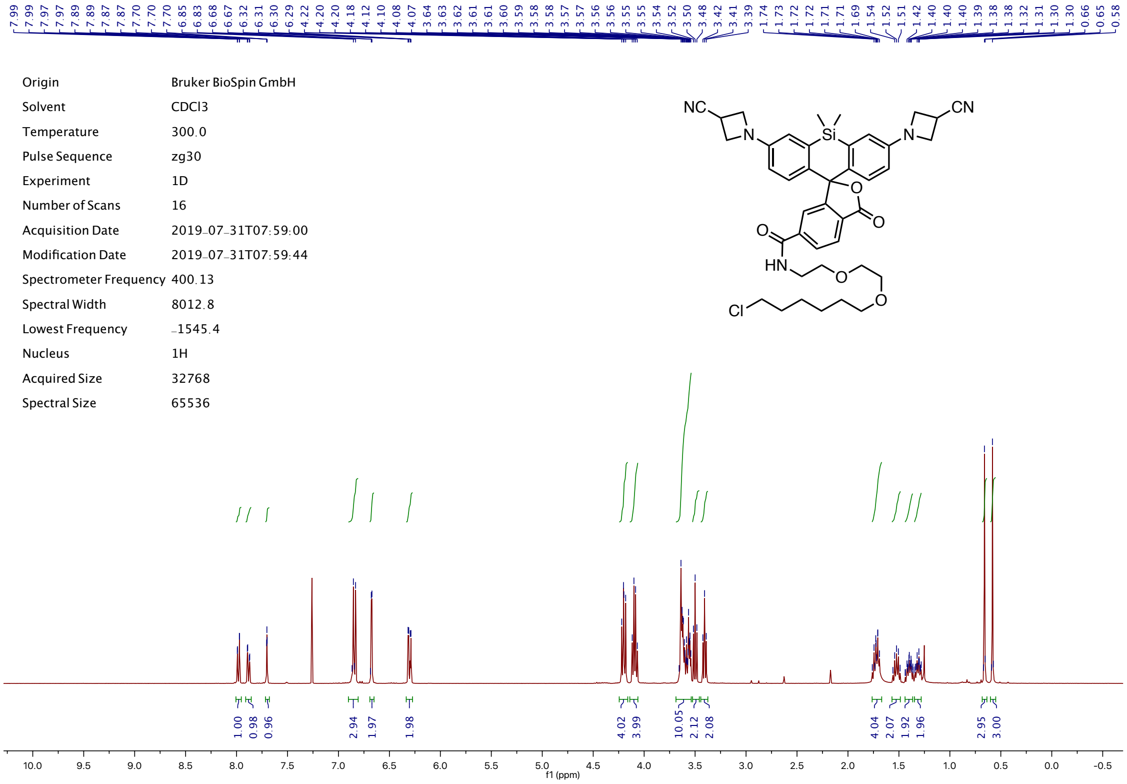
Origin Bruker BioSpin GmbH
Solvent CDCl3
Temperature 300.0
Pulse Sequence zg30
Experiment 1D
Number of Scans 16
Acquisition Date 2019-07-31T07:59:00
Modification Date 2019-07-31T07:59:44
Spectrometer Frequency 400.13
Spectral Width 8012.8
Lowest Frequency -1545.4
Nucleus 1H
Acquired Size 32768
Spectral Size 65536
1.00 0.98 0.96 2.94 1.97 1.98 4.02 3.99 10.05 2.12 2.08 4.04 2.07 1.92 1.96 2.95 3.00
f1 (ppm)

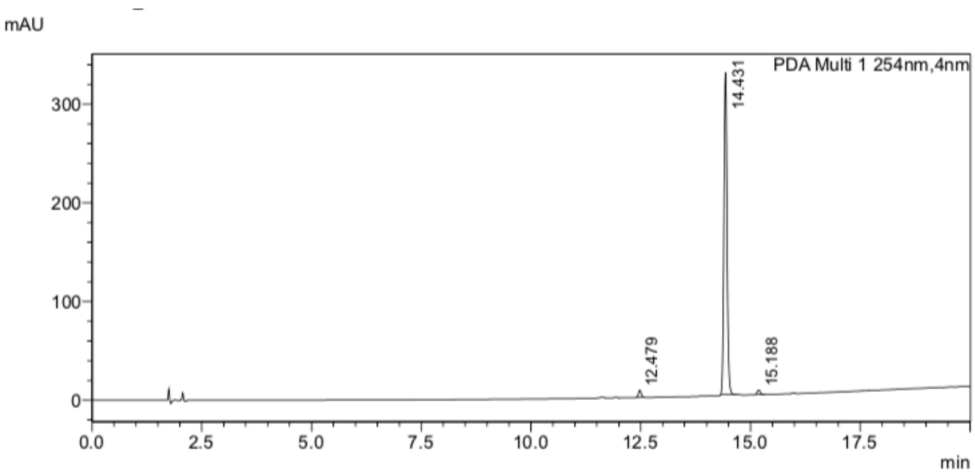
mAU
PDA Multi 1 254nm,4nm
14.431
12.479
15.188
min

| | |
|---|---|
| Origin | Bruker BioSpin GmbH |
| Solvent | CDCl3 |
| Temperature | 300.0 |
| Pulse Sequence | zg30 |
| Experiment | 1D |
| Number of Scans | 16 |
| Acquisition Date | 2019-07-31T07:54:00 |
| Modification Date | 2019-07-31T07:54:42 |
| Spectrometer Frequency | 400.13 |
| Spectral Width | 8012.8 |
| Lowest Frequency | -1545.4 |
| Nucleus | 1H |
| Acquired Size | 32768 |
| Spectral Size | 65536 |

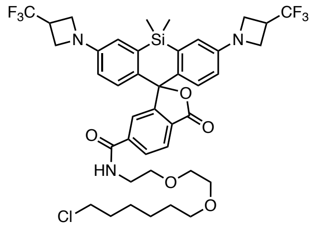


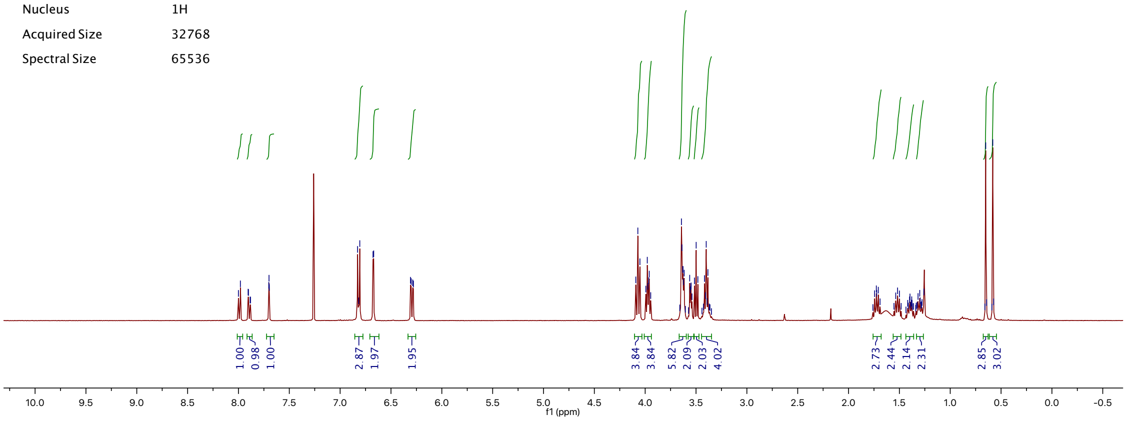


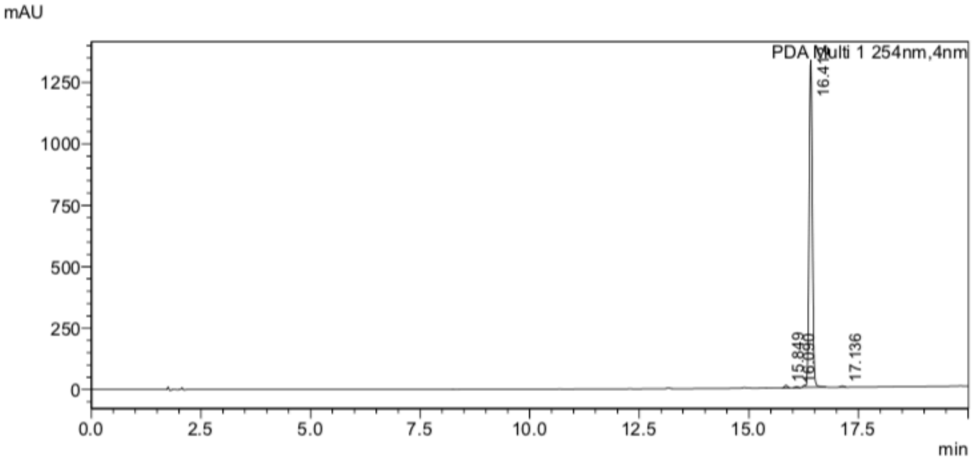

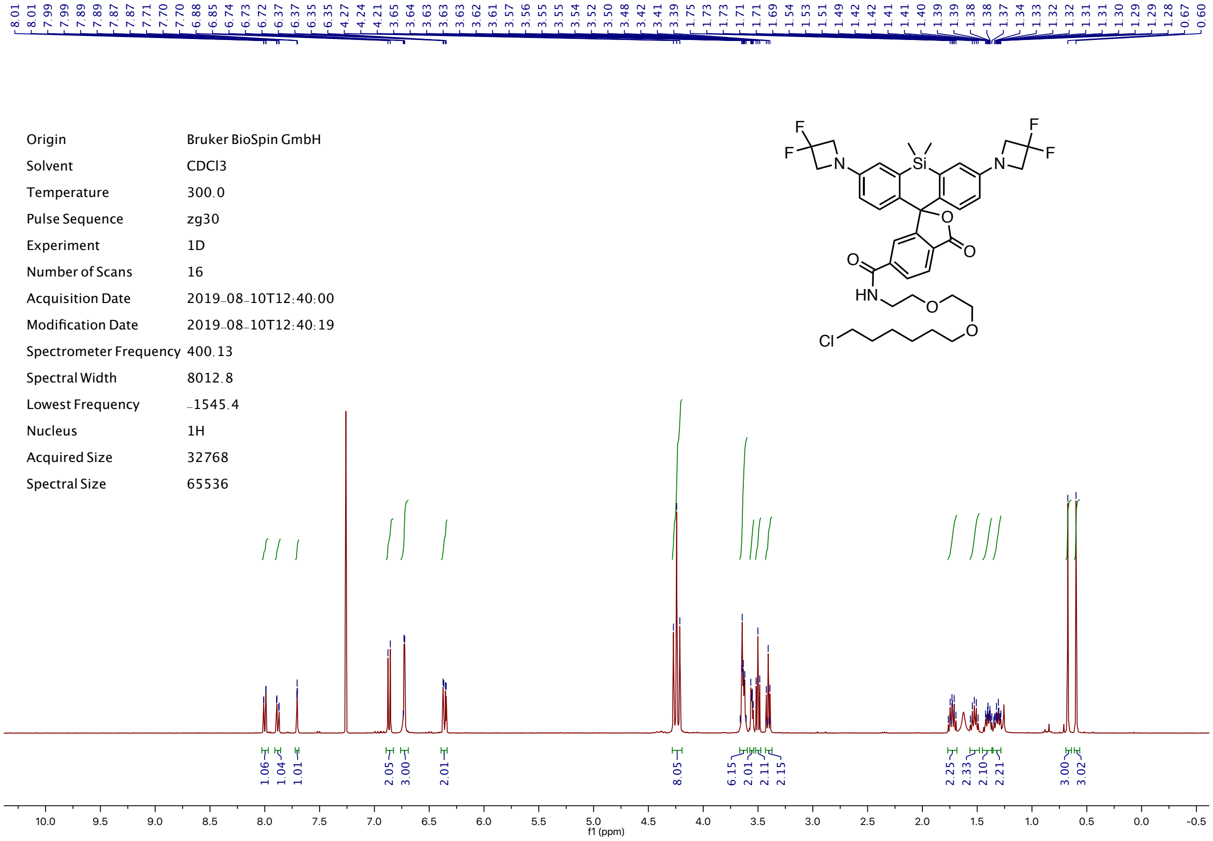
Origin Bruker BioSpin GmbH
Solvent CDCl3
Temperature 300.0
Pulse Sequence zg30
Experiment 1D
Number of Scans 16
Acquisition Date 2019-08-10T12:40:00
Modification Date 2019-08-10T12:40:19
Spectrometer Frequency 400.13
Spectral Width 8012.8
Lowest Frequency -1545.4
Nucleus 1H
Acquired Size 32768
Spectral Size 65536
f1 (ppm)

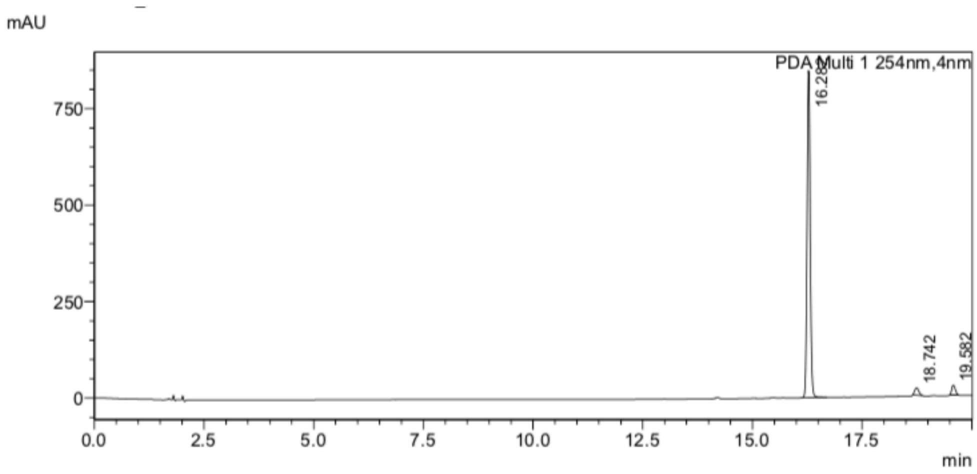
mAU
PDA Multi 1 254nm,4nm
18.742
19.582
min
